# Identification and characterization of a ubiquitin E3 RING ligase of the *Chlamydia*-like bacterium *Simkania negevensis*

**DOI:** 10.1101/2024.11.19.624306

**Authors:** Eva-Maria Hörner, Vanessa Boll, Thomas Hermanns, Adriana Moldovan, Kay Hofmann, Vera Kozjak-Pavlovic

**Affiliations:** Chair of Microbiology, Biocenter, Julius-Maximilian University Würzburg, D-97074 Würzburg, Germany; Institute for Genetics, University of Cologne, Zülpicher Straße 47a, D-50674 Cologne, Germany

**Author notes:** Corresponding author: Vera Kozjak-Pavlovic; Biocenter, Am Hubland, D-97074 Würzburg, Germany.

**Keywords:** RING ligase, Simkania negevensis, Chlamydia, ubiquitin, intracellular bacteria, infection

## Abstract

In the arms race between a pathogen and the host, many bacteria have acquired a sizable armory to counteract or change the defense mechanisms of the host cell, including the eukaryotic ubiquitin system. *Simkania negevensis*, an obligate intracellular *Chlamydia*-like bacterium, is one such example. *S. negevensis* has a biphasic developmental cycle, similar to other members of the *Chlamydiales* order. The bacterium develops inside a tubular membranous compartment called *Simkania*-containing vacuole, which is in close contact with the host endoplasmic reticulum (ER) and mitochondria. It can infect a wide range of hosts, with a long infection cycle lasting up to 15 days, and is associated with respiratory tract diseases. Recently, *S. negevensis* has been discovered to possess an unusually large number of deubiquitinating enzymes, but not much is known about the variety of other ubiquitin-modifying enzymes in this bacterium. Our study provides an initial description of the activity of a so far uncharacterized *S. negevensis* ubiquitin E3 RING-ligase (SNE_A12920 or SneRING). We report that SneRING primarily generates K63- and K-11 linked ubiquitin chains and preferentially interacts with UbcH5b and UBE2T E2 ubiquitin ligases among the ones we tested. Bacteria express SneRING upon infection of various human cell lines, as well as amoeba. In addition, we demonstrate that a portion of the expressed SneRING co-localizes with mitochondria and ER. Mass spectrometry studies of the SneRING interactome show enrichment in mitochondrial and ER proteins containing a prohibitin domain and involved in organelle morphology, respiration, and stress response. Our work offers the first insights into the function of an *S. negevensis* RING ligase, an enzyme potentially involved in organelle remodeling to accommodate the unique lifestyle of this intracellular bacterium.

## Introduction

*Simkania negevensis* (Sne) is an obligate intracellular *Chlamydia*-like bacterium, first described in 1993 as a cell culture contaminant of unknown origin (Kahane *et al*, 1993). Based on sequence comparison of the ribosomal DNA (rDNA), it has been classified as a member of the family *Simkaniaceae* within the order *Chlamydiales* (Everett *et al*, 1999). The bacterium survives and replicates in a great variety of hosts ranging from different eukaryotic cells including immune cells such as macrophages (Kahane *et al*, 2008; Kahane *et al*, 2007) to arthropods (Croxatto *et al*, 2014) and amoeba (Kahane *et al*, 2001). However, its natural host is unknown. Sne has been associated with different infections of the respiratory tract, including acute bronchiolitis in infants (Kahane *et al*, 1998) and community-acquired pneumonia (Lieberman *et al*, 1997). Furthermore, patients positive for Sne had a higher risk of transplanted lung rejection (Husain *et al*, 2007; Jamal *et al*, 2015). In addition, Sne has been also connected to Crohńs disease (Scaioli *et al*, 2019). However, recent studies dispute the link between Sne and respiratory tract infections (Al-Younes *et al*, 2017). While some of the closely related *Chlamydia*-species, such as Chlamydia trachomatis, the causative agent of sexually transmitted disease and trachoma (Newman *et al*, 2015), or Chlamydia pneumoniae, which causes acute respiratory diseases (Kuo *et al*, 1986), are highly pathogenic, it remains unclear to what extend Sne can cause infections in immunocompetent individuals.

Like other members of the *Chlamydiales*, Sne exhibits a biphasic developmental cycle, which is characterized by the alternating differentiation into two forms. While Reticulate Bodies (RB) represent the large, replicative, metabolically active intracellular form, which is stained homogeneously, the infective smaller and electron-dense form resembles the environmental Elementary Bodies (EB) of Chlamydia (Kahane *et al*., 1993; Kahane *et al*, 2002). Inside the host cell, Sne replicates within a Simkania-containing vacuole (SnCV). The SnCV is a membranous, tubular system that is associated with mitochondria and forms extensive contact sites with the endoplasmic reticulum (ER) of the host cell (Mehlitz *et al*, 2014). Sne exhibits a significantly longer life cycle than C. trachomatis. SnCV development and number of RBs plateau on day 3 post-infection (pi). An increased release of Sne infectious particles from infected HeLa or differentiated THP-1 cells starts on day 4 or day 5 pi, respectively (Koch *et al*, 2020), which is longer than the usual 2-3 days for C. trachomatis (Jury *et al*, 2023). In addition, Sne-infected cells can be observed in culture for up to 15 days (Kahane *et al*., 2002).

Given that bacteria proliferating within the host cell are exposed to the host cytoplasm, they risk being targeted by the cell-autonomous immunity. This includes the ubiquitination of free-living bacteria as well as bacteria that live in specialized vacuoles (Hermanns & Hofmann, 2019). Ubiquitination is a dynamic, versatile protein modification that regulates various cellular aspects in eukaryotes. The covalent attachment of ubiquitin to the targeted protein occurs through a cascade with three different enzymes: the E1 ubiquitin-activating enzyme, the E2 ubiquitin-conjugating enzyme, and the E3 ubiquitin-ligating enzyme. During ubiquitination, a single ubiquitin molecule is attached to a lysine residue of a specific substrate through an isopeptide bond. The complexity of ubiquitination is further increased by the property of ubiquitin itself to become ubiquitinated at one or more of its seven lysine residues, giving rise to ubiquitin chains. In addition, also the N-terminus of ubiquitin can be modified, which results in the possibility of forming eight different linkage types (Swatek & Komander, 2016).

Depending on the type of the ubiquitin chain, the outcome for the modified substrate differs. The prototypical ubiquitin chains are linked by lysine (K) 48 or 63 of the ubiquitin moieties. While K48- linked ubiquitin chains target their substrate for proteasomal degradation, K63-linked ubiquitin chains regulate NF-kB transcription factor activation, responses of the immune system, repair of DNA, sorting of proteins, clearance of damaged mitochondria and they can also control the assembly of large protein complexes that drive translation or splicing of mRNA. Ubiquitin chains, connected via the free N-terminus or the remaining five lysine residues are said to be atypical conjugates. These include K11-, K6-, and M1-linked ubiquitin chains. K-11 linked chains play a role during cell division by targeting cell cycle regulators for degradation. Interestingly, K11-conjugates are also associated with innate immune response against viruses (Yau & Rape, 2016). Although not much is known about the cellular function of K6-linked ubiquitin chains, they have been observed to accumulate upon depolarization of mitochondria and after UV radiation. Ubiquitin moieties linked via their free N-terminus, also referred to as linear or M1-linked ubiquitin chains, are involved in the regulation of immune response and inflammation through regulating the activation of the NF-kB transcription factor (Akutsu *et al*, 2016).

A recent discovery of branched and mixed ubiquitin chains forms another level of ubiquitination diversity. Determining the function of branched polymers is challenging but gives new insights into the variety of information that can be transported by ubiquitin signals. K11/K48 heterotypic ubiquitin chains are assembled during mitosis and on misfolded proteins in the ER or cytoplasm. They have been shown to play a role in cell cycle control, enhance substrate recognition, and accelerate proteasomal degradation. In contrast, branched K48/K63 conjugates are synthesized in response to the activation of the transcription factor NF-kB. Additionally, modification of substrates during apoptotic response with this mixed chain type results in their degradation by the proteasome (French *et al*, 2021; Haakonsen & Rape, 2019). For mixed K11/K63-linked polymers, a non-proteolytic function during endocytosis and signaling via the transcription factor NF-kB has been determined (Wickliffe *et al*, 2011). Even though K6/K11 and K6/K48 branched ubiquitin chains have already been identified, their functions remain unclear (Haakonsen & Rape, 2019).

Based on their structure and mechanism of ubiquitin transfer, E3 ubiquitin ligases are usually divided into three classes: RING (really interesting new gene), RBR (RING-between-RING), and HECT (homologous to the E6/AP carboxyl terminus) ligases. While RBR and HECT E3s form a catalytic intermediate with ubiquitin by binding the ubiquitin from the loaded E2 enzyme to themselves before conjugating it to the target protein, RING ligases directly transfer ubiquitin from the E2 enzyme to the substrate (Toma-Fukai & Shimizu, 2021). With more than 600 RING ligases encoded by the human genome, this group is the largest. RING E3s are characterized by a RING domain that directly binds the substrate and the ubiquitin-loaded E2 enzyme. By bringing both components into proximity, RING ligases catalyze ubiquitin transfer from the conjugating enzyme to the target protein (Deshaies & Joazeiro, 2009; Zheng & Shabek, 2017).

Many pathogenic bacteria have acquired a sophisticated range of effectors, including ubiquitin E3 ligases, to exploit the host cell’s ubiquitin system for their benefit. Examples include SopA, a HECT-like E3 ligase from Salmonella, LubX, a RING-like E3 ligase from Legionella, as well as several novel E3 ligases from Shigella (Maculins *et al*, 2016). By ubiquitinating the substrates, they modify the ubiquitin code to form a niche that allows them to survive and multiply within the host cell (Dikic & Schulman, 2023).

Since Sne is capable of long infections and intracellular growth while simultaneously efficiently suppressing the host defense system, it is of great interest how the bacterium accomplishes this. Recent studies showed that Sne possesses an unusually high number of deubiquitinating enzymes (DUBs) with sometimes interesting cleavage specificities. Furthermore, these DUBs are from diverse classes, of which some have never been described in bacteria (Boll *et al*, 2023).

In this work, we focused on identifying and characterizing putative ubiquitin (E3) RING ligases of Sne. We focused on a RING ligase Sne_A12920 (SneRING), a 25.4 kDa protein containing an unusual RING finger when compared to other bacterial RING ligases. We purified the enzyme and determined its ubiquitinating activity, as well as detected the region of interaction with the E2 ubiquitin-conjugating enzyme. Additionally, we demonstrated that the SneRING shows different ubiquitination activity depending on the E2 conjugating enzyme it interacts with. Finally, we determined that the ligase generates specific types of ubiquitin chains, rather than producing all chain types. SneRING is expressed in different human cell lines as well as in amoeba during infection. After expression in human cells, a portion of the protein is associated with the ER and mitochondria. The possible interaction partners of SneRING include mitochondrial and ER proteins that play a role in morphology, energetics, and transport, indicating the potential mode of action of this bacterial enzyme. Taken together, our data provide first insights into the previously unknown world of ubiquitin ligases in *Simkania negevensis*.

## Results

### Identification of RING ligases in *Simkania negevensis*

To identify ubiquitin ligase candidates in the sequenced genome of Sne (Collingro *et al*, 2011), the Sne subset of UniProt (UniProtConsortium, 2023) was searched with generalized profiles (Bucher *et al*, 1996) representing classical RING finger domains and several RING finger variants. When using a profile derived from a multiple alignment of established RING domains, only a single Sne ORF, Sne_A12920 (SneRING), reached a significant score of p<0.01. To validate this finding, BLAST and profile searches were used to find other bacterial proteins related to SneRING. A small family was identified, of which representative members include Sim-KFB93 and Sim-QNJ27 from bacteria belonging to the *Simkaniaceae* family, and Chl-HKST.2, belonging to an unclassified *Chlamydiia* (Fig 1A). An Alphafold (Jumper *et al*, 2021) model of SneRING shows a two-domain architecture with a RING-like fold forming the first domain (residues 14-72), followed by a linker region and a C-terminal α/β-fold domain without informative sequence or structural similarities (Fig 1B). Like canonical RING fingers, the RING-like domain of SneRING is predicted to coordinate two Zn^2+^ ions, albeit using somewhat atypical ligand residues. In the model, the first Zn^2+^ ion is coordinated by Cys-17, Cys-20, His-40, and Cys43, while the second Zn^2+^ ion is contacted by Asn-32, Cys-35, Cys-54, and Cys57.

**Figure 1.**
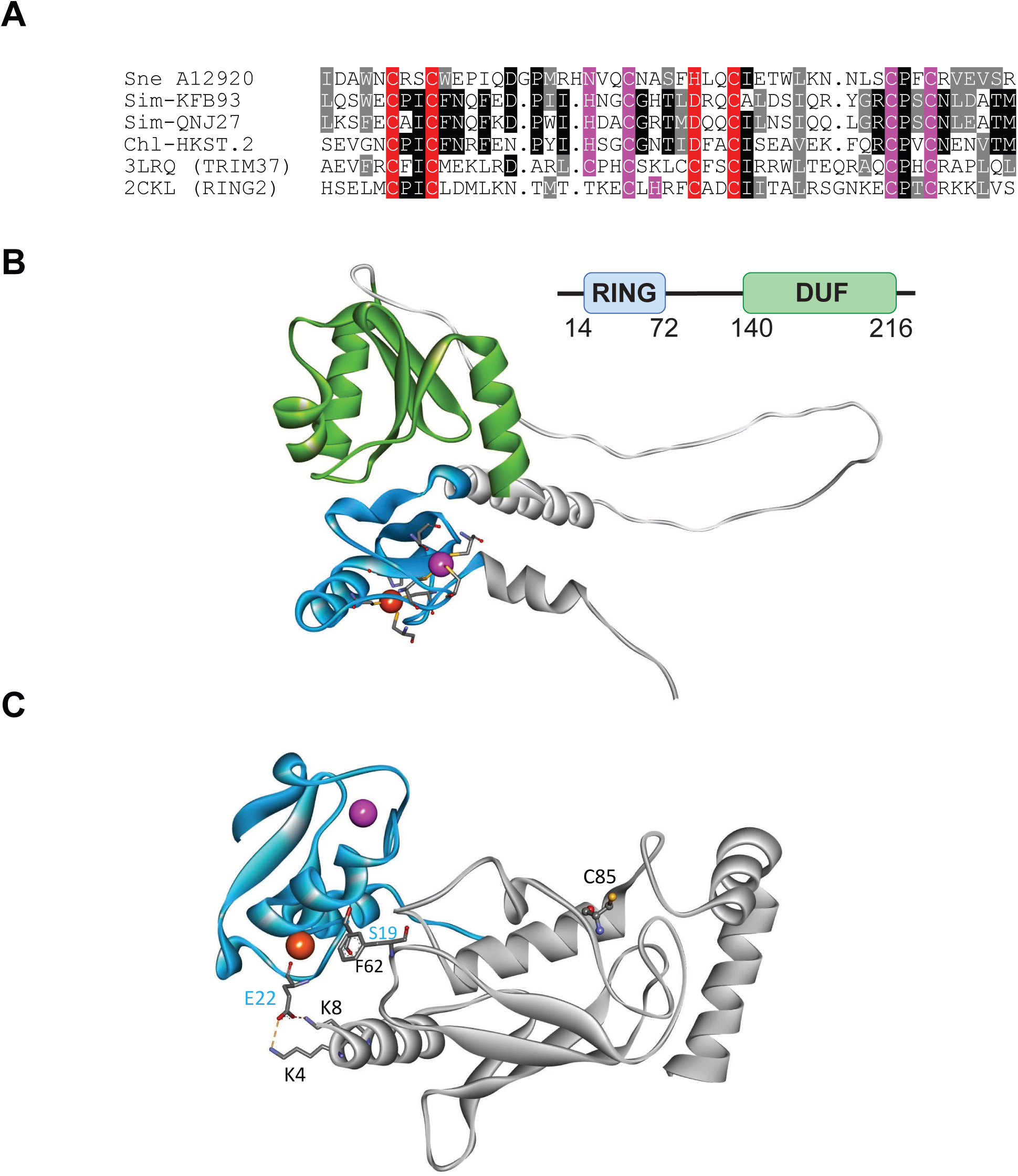
SneRING is a predicted RING domain ligase. **(A)** Multiple alignment of the RING domains of SneRING (Sne_A12920), some bacterial relatives (Sim-KFB93: *Simkaniaceae* bacterium RefSeq:QVL56959; Sim-QNJ27: *Simkaniaceae* bacterium RefSeq:MDJ0651792; Chl-HKST: *Chlamydiia* bacterium Uniprot: A0A960X5Q0), and sequences of the two best DALI hits (pdb:3LRQ (TRIM37 from *Homo sapiens*) and pdb:2CKL (RING2 from *Mus musculus*)). Residues invariant or conserved in at least 50% of the sequences are shown on black and grey background, respectively. Residues involved in the coordination of Zn_1_ and Zn_2_ are highlighted in red and magenta, respectively. **(B)** Alphafold model of SneRING shows the N-terminal RING-like domain (blue) and a C-terminal domain of unknown function (DUF, green). The Zn_1_ and Zn_2_ ions coordinated by the RING-like domain are shown in red and magenta, respectively. **(C)** Alphafold model of the RING domain of SneRING (blue) in contact with UbcH5b (grey). The Zn_1_ and Zn_2_ ions coordinated by the RING-like domain are shown in red and magenta, respectively. The predicted contact residues Glu-22 (E22) and Ser-19 (S19) and their interaction partners are labeled.

A DALI (Holm, 2022) search using the RING-like region of SneRING identified as significant top-hits the established RING fingers of TRIM37 (pdb:3LRQ, Z-score 9.1) and RING2 (pdb:6WI7, Z-score 8.9). RING2 possesses a typical RING finger, where Zn_1_ is coordinated by two CxxC dyads and Zn_2_ is bound by a CxH and a CxxC dyad. In SneRING, both coordination spheres are slightly altered: His-40 replaces Cys as the 3^rd^ ligand of Zn_1_, while the first two ligands of Zn_2_are NxxC rather than the canonical CxH (Fig 1A). Functionally, these changes might be inconsequential, as Alphafold model of the Sne_A12920 RING domain in contact with UbcH5 predicts SneRING to form a complex with E2 enzymes resembling that of other RING ligases (Fig 1C). As with other RING fingers, the most important contacts are made by the region surrounding the first Zn-binding dyad. While most other RING domains contact E2s by a conserved hydrophobic residue before the 2^nd^ Zn-binding cysteine, SneRING carries the atypical Ser-19 at this position, which does not form a specific E2 interaction in the model. Instead, Glu-22 forms short-distance salt bridges to Lys-4 and Lys-8 of UbcH5 (Fig 1C). We reasoned that a mutation of Glu-22 combined with a Ser-19 mutant might abrogate E2-binding. Indeed, when subjecting an S19R_E22R variant of SneRING to Alphafold modeling with UbcH5, no productive complex was formed.

Besides the significantly-scoring SneRING, three further RING-like domains could be identified bioinformatically. A closer inspection of the borderline-significant matches identified the uncharacterized ORF Sne_A08700 and its close relative (and genomic neighbor) Sne_A08690 to have a conspicuous arrangement of cysteine and histidine residues. The assumed RING-relationship of these two proteins is supported by the conservation pattern of their sequence family and by Alphafold modeling: Sne_A08700 is predicted to use eight canonical ligands for coordinating two Zn^2+^ions (Fig S1A); the best DALI match to an established RING-finger was RNF125 (pdb:5DKA, Z-score 5.6). Sequence conservation between RNF125 and the Sne_A08700 family shows a big insertion after the 3^rd^ Zn ligand (Fig S1B), which explains the poor score in the initial profile search. The closely related protein Sne_A08690 has lost three ligands for the first Zn^2+^ ion and probably coordinates only a single ion. A 4^th^ RING candidate (Sne_A19470) was picked up in profile searches using the SP-RING family, a subgroup of RING-like domains usually involved in SUMO conjugation (Hochstrasser, 2001). However, a closer inspection suggested that Sne_A19470 does not contain an SP-RING domain but rather a (degenerate) conventional RING that has lost its ligands for Zn_1_ coordination (Fig S1C). The best DALI matches were TRIM69 (pdb:6YXE, Z-score 8.5) and TRIM2 (pdb:8A38, Z-score 8.3), which both contain a structurally similar RING domain, but with a complete set of eight ligand residues (Fig S1D).

Considering that SneRING was the only protein with a RING-like domain that has been identified in a previously published proteomics analysis of the ER/SnCV membrane of infected HeLa229 cells on day 3 pi (Herweg *et al*, 2016), we focused on further characterization of this enzyme.

### SneRING is an active E3 RING ligase with an E2 enzyme-interacting region within its predicted RING domain

Our bioinformatic analysis only predicts putative RING ligases without hinting at the protein function. Therefore, we first tested the activity of the SneRING by analyzing ubiquitin chain assembly during an *in vitro* autoubiquitination assay. To do so, we purified SneRING and incubated it with a ubiquitin-activating (E1) and a ubiquitin-conjugating enzyme (E2) as well as free ubiquitin and ATP to allow the enzyme cascade to covalently attach ubiquitin to SneRING. Already after 30 min of incubation, we could detect a strong ubiquitin signal that increased over time, while in the negative control without SneRING, only monoubiquitin could be detected (Fig 2A). This indicates that the SneRING is an active ubiquitin ligase.

**Figure 2.**
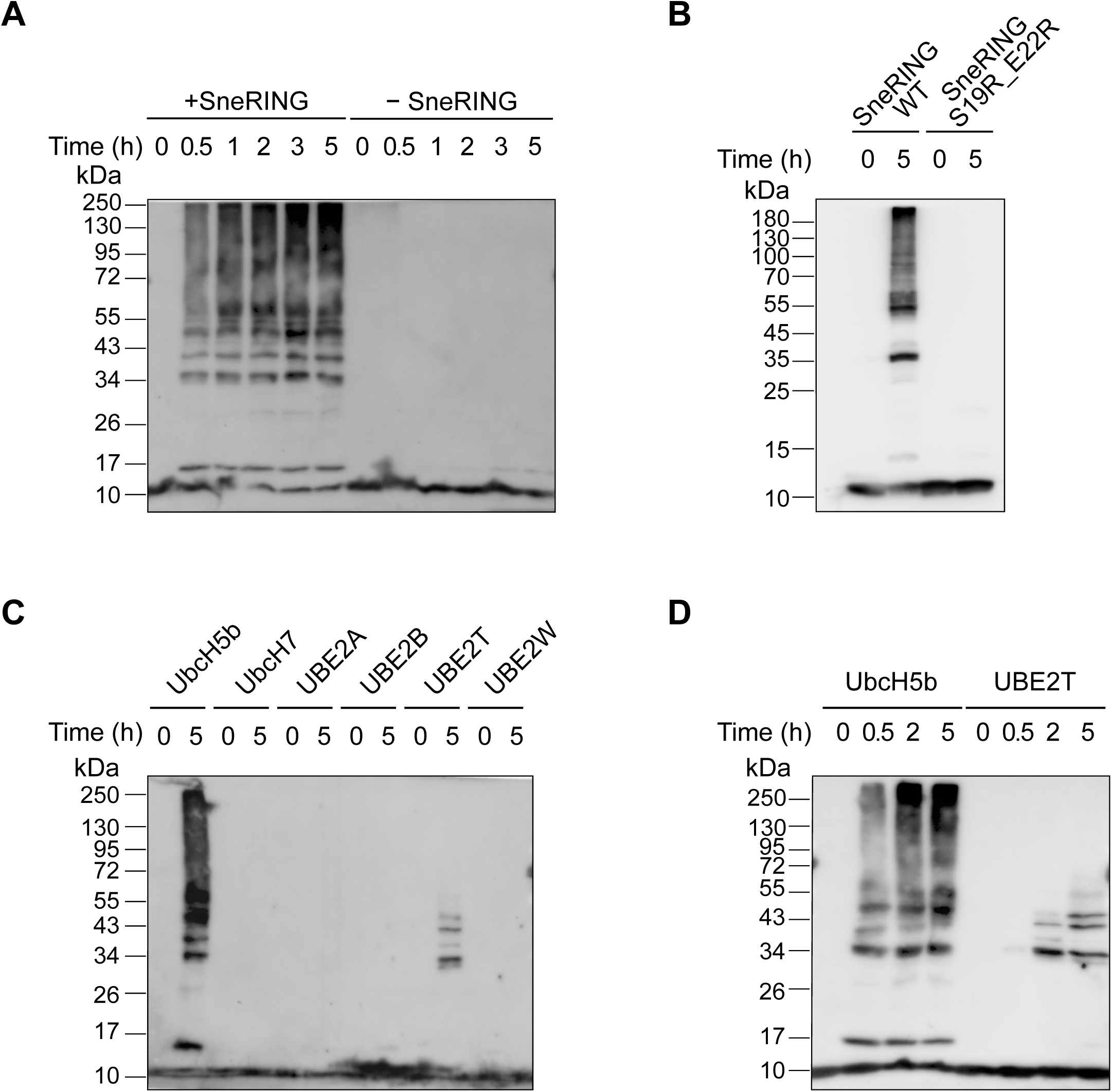
SneRING shows ubiquitination activity in an *in vitro* assay. **(A)** Purified recombinant SneRING was mixed with recombinant E1 ubiquitin-activating enzyme, E2 ubiquitin-conjugating enzyme (UbcH5b), ubiquitin, and ATP for indicated periods. A reaction without the ligase served as a control. Samples were analyzed by SDS-PAGE and western blot, using an antibody against ubiquitin. **(B)** *in vitro* autoubiquitination reaction of wildtype SneRING and SneRING-ligase with mutated E2 enzyme interacting region (SneRING S19R_E22R) was performed for 5 h as in A and analyzed by SDS-PAGE and western blot, using anti-ubiquitin antibodies. **(C)** *in vitro* autoubiquitination reaction of wildtype SneRING was performed as in A in combination with indicated E2 enzymes for 5 h, followed by SDS-PAGE and western blot analysis using ubiquitin antibody. **(D)** SneRING was incubated with E2 enzyme UbcH5b or UBE2T in an *in vitro* autoubiquitination assay as described in A. Samples were taken at indicated time points and tested by immunoblot using the primary antibody against ubiquitin.

Since Alphafold modeling indicated the importance of Glu-22 and Ser-19 for the interaction with an E2 enzyme, we replaced these two amino acids with arginine to generate a catalytically inactive form and performed an *in vitro* autoubiquitination assay with the purified mutated enzyme for 5 h. While in the presence of the wild-type SneRING a strong ubiquitin signal could be detected, no ubiquitin chains were formed in the reaction containing SneRING S19R_E22R (Fig 2B). This confirms that the N-terminal domain of the SneRING interacts with the E2 ubiquitin-conjugating enzyme and is necessary for the enzyme function.

To test whether SneRING can generate ubiquitin chains when combined with other E2 enzymes besides UbcH5b, we performed an *in vitro* autoubiquitination assay for 5 h using selected ubiquitin-conjugating enzymes. Only UbcH5b- and, to a lesser extent, UBE2T-containing reactions showed the presence of the poly-ubiquitin signals, while no ubiquitin chain assembly occurred with the addition of the E2 enzymes UbcH7, UBE2A, and UBE2W (Fig 2C). The time course analysis of the autoubiquitination reaction in combination with UbcH5b showed strong ubiquitination after 30 min of incubation, increasing over time (Fig 2A,D). In contrast, in the presence of UBE2T nearly no reaction was observed after 30 min, while 2 h and 5 h later, some poly-ubiquitin signal was detectable (Fig 2D). Among the tested E2 enzymes, SneRING appears to be most active in combination with UbcH5b, while a moderate reaction also occurs together with the ubiquitin-conjugating enzyme UBE2T.

### Ubiquitin chains generated by SneRING are cleaved by deubiquitinating enzymes specific for the K11/K63 linkage type

To characterize the type of ubiquitin chains formed by SneRING, we performed linkage-specific chain-cleaving assays (UbiCRest assays) (Hospenthal *et al*, 2015). Ubiquitin chains generated by SneRING after *in vitro* autoubiquitination for 5 h were incubated with a panel of different DUBs that either process any chain type, such as ubiquitin-specific peptidase 21 (USP21) (Ye *et al*, 2011), or have specificity against certain ubiquitin chain linkages. When incubating the reaction with the USP21, no remaining polyubiquitin signal could be detected. Treatment with the K11-specific DUB Cezanne (Bremm *et al*, 2010) or with the K63-specific DUB AMSH (associated molecule with an Src homology 3 domain of signal transducing adaptor molecule, STAM) (Komander *et al*, 2009), strongly reduced the polyubiquitin signal. In contrast, no cleavage was observed upon incubation with the DUB SneOTU that specifically cleaves linear M1 ubiquitin chains or SneVTD that shows cleavage specificity against K6-linked ubiquitin chains (Boll *et al*., 2023). Furthermore, also the K48-specific DUB OTUB1 (Wang *et al*, 2009) did not visibly reduce the ubiquitin signal (Fig 3A).

**Figure 3.**
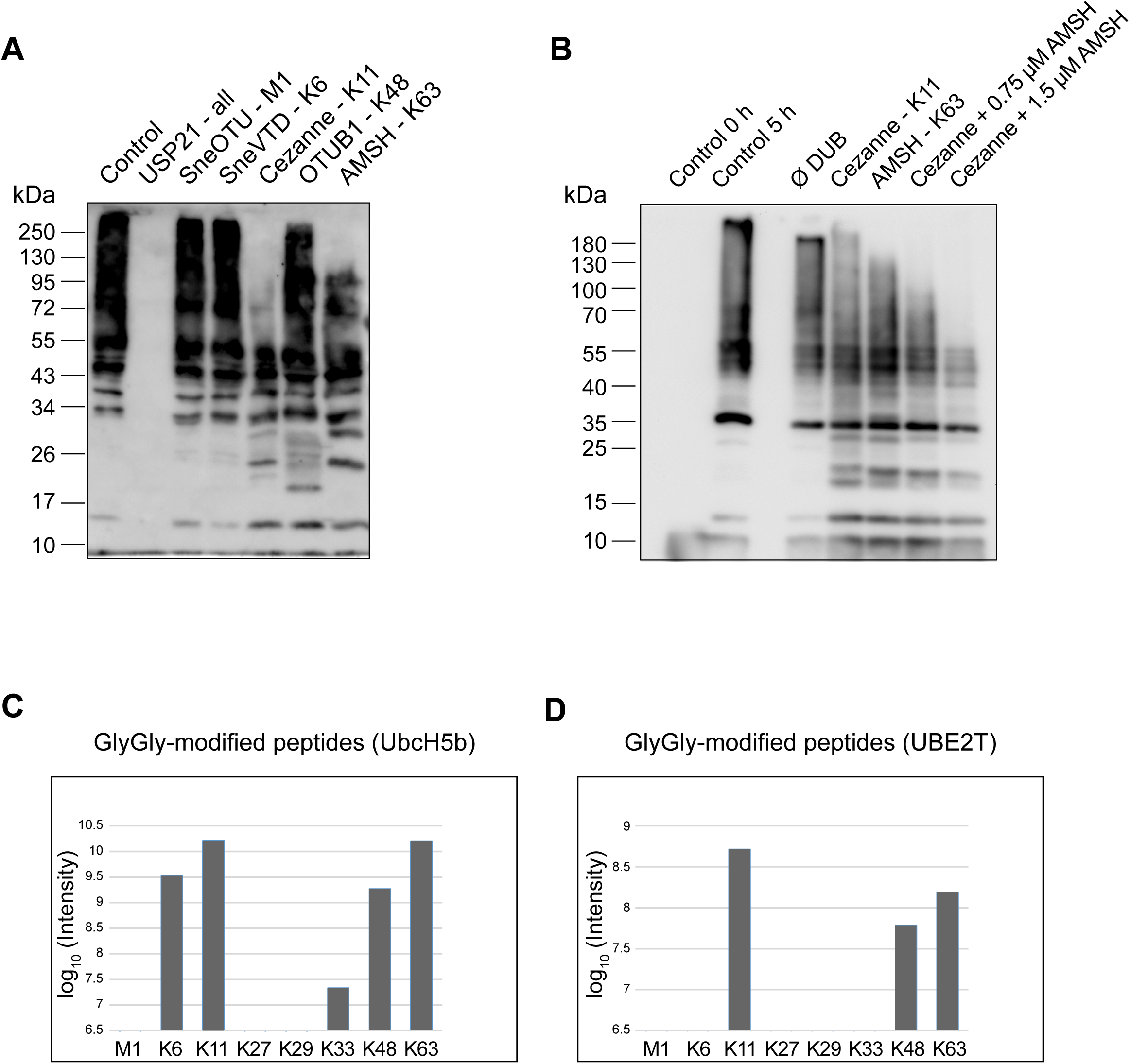
SneRING generates K6-, K11-, K48, and K63-linked ubiquitin chains that can be digested by specific DUBs. **(A)** An *in vitro* autoubiquitination reaction as in Fig. 2A was incubated for 5 h, after which an UbiCRest assay was performed using the indicated chain type-specific DUBs for 1 h at room temperature. Ubiquitin chain cleavage was analyzed by SDS-PAGE and western blot using a primary antibody against ubiquitin. **(B)** An *in vitro* autoubiquitination reaction as in A was incubated with Cezanne (K11-linked ubiquitin chain-specific DUB) and AMSH (K63-linked ubiquitin chain-specific DUB). Degradation of ubiquitin chains was detected by SDS-PAGE and western blot using anti-ubiquitin antibody. **(C, D)** A 5 h *in vitro* autoubiquitination reaction of SneRING with UbcH5b (C) or UBE2T (D) was analyzed by mass spectrometry. The graphs show log_10_ intensities of GlyGly-modified residues for the different linkage types, as determined by MaxQuant.

To test whether the ligase only generates these two linkage types, or if other ubiquitin chain types are produced at a lower level, a di-UbiCRest assay was performed. To do so, next to incubating an *in vitro* SneRING autoubiquitination reaction with the K11- specific DUB Cezanne or with the K63- specific DUB AMSH, the reaction was also incubated with both enzymes simultaneously. As already observed, Cezanne and AMSH led to a reduction of the ubiquitin signal, demonstrating the presence of K11- and K63-linked ubiquitin chains. However, the combined activity of both DUBs did not completely degrade the ubiquitin chains generated by the SneRING (Fig 3B), indicating that other linkage types are formed by the SneRING in addition to K11- and K63-linked ubiquitin chains. Mass spectrometry of the *in vitro* SneRING autoubiquitination reaction in the presence of UbcH5b (UBE2D2) mainly detected K11- and K63-linked GlyGly-modified peptides, with some signals for K6- and K48-modified peptides also visible. Similar results with overall weaker signal strength were obtained for the reaction containing the E2 enzyme UBE2T (Fig 3C). In conclusion, SneRING forms *in vitro* mostly K11- and K63-, in addition to K6- and K48-linked ubiquitin chains.

To further validate this finding, *in vitro* autoubiquitination assays performed with UbcH5b were tested by immunoblot analysis for the presence of specific ubiquitin chains. We detected strong signals using antibodies against K11- (Fig S2A) and K63-linked chains (Fig S2B). For K6- and K48-linked chains we used native linked di-Ubiquitin conjugates as positive controls (UbiQ). The antibody against K48-linked ubiquitin chains produced a signal in the control sample, as well as in the ubiquitin chains generated by SneRING (Fig S2C). Similar observations were made when using an antibody against K6-linked ubiquitin conjugates (Fig S2D). However, when we used an antibody against M1- linkages, which were not detected by UbiCRest assay and mass spectrometry, we only saw a signal in the positive control, but not in the SneRING reaction, confirming the previous results (Fig S2E).

### SneRING is expressed during infection of human cells and its possible natural host Acanthamoeba castellanii

We next analyzed the expression pattern of SneRING during infection. Since Sne can infect and grow in a variety of cells (Vouga *et al*, 2017), as well as survive and multiply within Acanthamoeba castellanii, we measured the SneRING mRNA levels during the infection cycle of various cell lines and amoeba by RT-qPCR analysis and normalized it against the mRNA level of the bacterial 5S RNA. In comparison, we measured the mRNA levels of a highly-expressed gene, Sne chaperone SnGroEL. SneRING was expressed in all host cells we tested, though to a lesser extent than SnGroEL (Fig S3). Expression intensities of SneRING in different human cell lines were comparable. In HeLa229 and U2OS cells we observed somewhat higher SneRING expression on day 2 pi compared to day 4 pi (Fig S3A,B).

Sne can infect and grow in macrophages (Kahane *et al*., 2008), as well as in the THP-1 cell line that mimics the macrophage-like state after differentiation by PMA (Chanput *et al*, 2014), in contrast to C. trachomatis (Herweg & Rudel, 2016). We analyzed and compared the expression of SneRING in the THP-1 cell line and primary human M2-like macrophages. Based on the observation that infective Sne particles can be detected on day 6 pi in macrophages (Kahane *et al*., 2008), we included this time point in our experimental setup. RT-qPCR analysis showed slightly higher expression of SneRING in THP-1 cells than in M2-like macrophages. There were no notable differences in the mRNA levels between different days pi (Fig S3C,D).

By taking up a high number of bacteria, many free-living protozoa mimic the role of professional phagocytes (Fritsche *et al*, 2000). Sne can survive and grow within amoeba (Kahane *et al*., 2001). Since Sne establishes a long-term infection in amoeba, we measured the mRNA level of SneRING only on day 4 pi. A. castellanii culture was cultivated at 30 °C instead of 37 °C that we used for cultivation of human cell lines (or 35 °C for Sne infected). mRNA of SneRING was detectable, but the levels varied strongly between the samples both for SneRING and SnGroEL (Fig S3E).

### SneRING localizes to host cell cytoplasm, mitochondria, and ER after overexpression, associating with the inner mitochondrial membrane

We next studied the localization of SneRING in the host cell. Since we were unsuccessful in producing an antibody against SneRING that would recognize the native protein, we limited our studies to the FLAG-tagged protein we expressed from a plasmid in U2OS cells. To increase the efficacy of SneRING expression, we used a human codon-adjusted version of the gene. SneRING was successfully expressed and exhibited cytosolic distribution as observed by immunofluorescence microscopy (Fig 4A). The FLAG-tagged protein could also be identified by western blot after transfection (Fig 4B). Upon subcellular fractionation and separation of the cells into the crude mitochondrial, the light membrane, and the cytosolic fraction, SneRING signal was distributed between all fractions (Fig 4C). In comparison, Mic60, a mitochondrial inner membrane protein, localized almost exclusively to mitochondrial fraction, while calnexin, the integral protein of the ER, was detected in both the light membrane and the crude mitochondrial fraction. GAPDH and tubulin were associated with every fraction but mainly localized in the cytosol, somewhat resembling SneRING distribution (Fig 4C).

**Figure 4.**
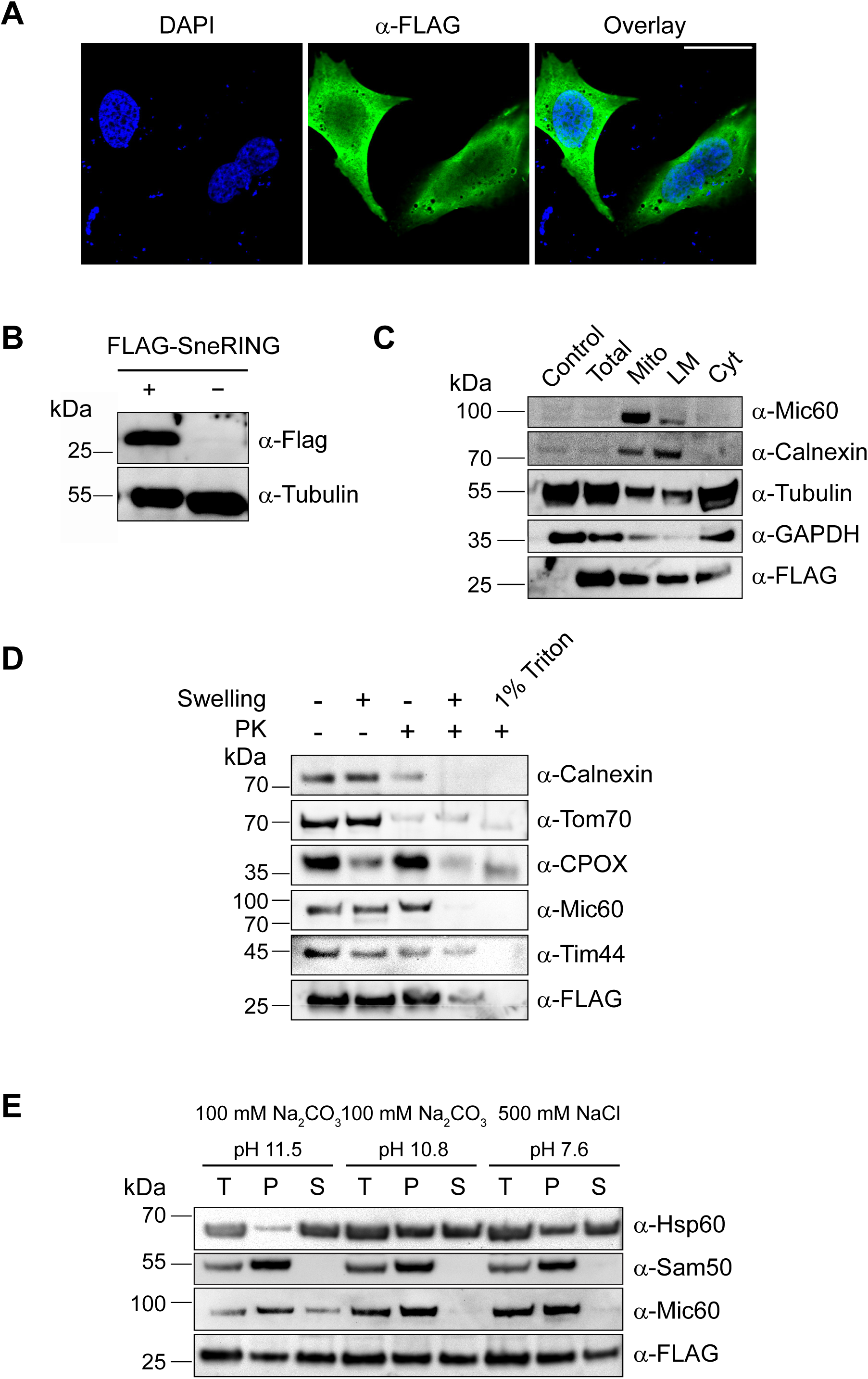
SneRING localizes to the host cell cytosolic, light membranes, and inner mitochondrial membrane after overexpression. **(A)** U2OS cells were transfected with a human codon-adjusted FLAG-tagged version of the SneRING using the transfection reagent PEI MAX®. 72 h post-transfection, cells were fixed and stained using DAPI (blue channel), and a primary antibody against FLAG followed by the fluorophore-coupled secondary antibody (green channel). Images were taken using laser scanning confocal microscopy. The scale bar represents 30 µm. **(B)** U2OS cells were transfected as in A, and 36 h post-transfection, analyzed by SDS-PAGE and western blot using antibodies against the FLAG-tag and tubulin. **(C)** U2OS cells were transfected as in A, and 48 h post-transfection, the cells were separated by differential centrifugation into a crude mitochondrial (Mito), light membrane (LM), and cytosolic (Cyt) fraction. Total cell lysate of non-transfected cells (Control) and transfected cells (Total) served as control. The samples were analyzed by SDS-PAGE and western blot using antibodies against Mic60, Calnexin, Tubulin, GAPDH, and FLAG. **(D)** U2OS cells were transfected as in D. After 48 h of expression, mitochondria were isolated and then incubated in isotonic buffer (-Swelling) or in hypotonic buffer (+Swelling) to rupture the outer mitochondrial membrane, combined with the treatment using 50 µg/mL of protease K (PK). For complete solubilization, mitochondria were treated with 1% Triton X-100 (1% Triton) and PK and precipitated using 72% trichloroacetic acid. Samples were analyzed by SDS-PAGE and western blot, using antibodies against Calnexin, Tom70, CPOX, Mic60, Tim44, and FLAG-tag. **(E)** Mitochondria as in D were either sonicated in a high-salt buffer (500 mM NaCl pH 7.6) or subjected to carbonate extraction (100 mM Na_2_CO_3_ pH 10.8 or 100 mM Na_2_CO_3_pH 11.5). Pellet (P) and supernatant (S) fractions were separated by ultracentrifugation, while untreated total samples (T) served as control. Samples were analyzed by SDS-PAGE and western blot using antibodies against Hsp60, Sam50, Mic60, and FLAG.

Immunofluorescence microscopy of HeLa229 cells with GFP-labeled mitochondria (Fig S4A) or dsRED-labeled ER (Fig S4B) that were transfected with FLAG-tagged SneRING gave a similar picture, with some overlap of the signal, indicating possible association of SneRING with these organelles. This did not significantly change upon infection with Sne (Fig S4A,B, lower panels). Notably, Sne-infected cells show fragmentation of the mitochondria, which is absent in cells transfected only with SneRING. In both cell lines, the FLAG-tagged protein is pushed to the cell periphery with the cytosol and is not identified in the SnCV when the cells are infected (Fig S4).

The observed association of SneRING with mitochondria could be caused by the interaction with the mitochondrial surface or result from the insufficient purity of the mitochondrial fraction, which contains ER and contaminations from other cellular compartments. To analyze this further, we isolated mitochondria from the FLAG-SneRING-transfected U2OS cells and subjected them to swelling in a hypotonic buffer in the absence or the presence of Protease K (PK). The accessibility of the proteins to PK demonstrates their submitochondrial localization. As a positive control, mitochondria were completely solubilized by 1% Triton X-100 and treated with PK (Fig 4D). Surprisingly, FLAG-tagged SneRING could be detected even after PK treatment of intact mitochondria, indicating that the protein is not localized on the mitochondrial surface. In contrast, Tom70, a mitochondrial outer membrane protein, is degraded under similar conditions, comparable to the ER protein calnexin. Opening of the outer membrane by swelling leads to a partial release of an intermembrane-space protein coproporphyrinogen oxidase (CPOX), but the inner membrane protein Mic60, the matrix protein Tim44, and SneRING remain undisturbed. PK treatment of the mitochondria with the ruptured outer membrane leads to a degradation of CPOX as well as Mic60, which is exposed to the intermembrane space, while Tim44 is degraded completely only after complete solubilization of mitochondria by a detergent. SneRING behaves in this respect like Mic60 since most of the protein is degraded upon PK treatment of swollen mitochondria. A small portion of the protein, however, remains undigested, indicating possible protection by the inner mitochondrial membrane (Fig 4D).

We performed carbonate extraction of isolated mitochondria using 100 mM Na_2_CO_3_, pH 11.5 or pH 10.8 to assess membrane integration of SneRING. In addition, the membrane association of the protein was tested by sonicating mitochondria in a high salt buffer. Hsp60, a soluble mitochondrial matrix protein, was found in both pellet and supernatant fractions after milder carbonate extraction and extraction in the high-salt buffer but was fully extracted at pH 11.5. In contrast, Sam50, an integral protein of the outer mitochondrial membrane, was only detected in the pellet after treatment in all three conditions. Mic60, the transmembrane protein of the inner mitochondrial membrane, was partially extracted from membranes only after carbonate extraction at the higher pH of 11.5. FLAG-tagged SneRING was detected in both the pellet and the supernatant after incubation in all three conditions, suggesting that the protein is mostly membrane-associated. A small portion, however, appears to be a membrane-integral protein (Fig 4E). We conclude that a portion of SneRING localizes to the host cell mitochondria after overexpression. The protein is most likely associated with the inner mitochondrial membrane.

### Putative SneRING-ligase interacting partners include mitochondrial and ER proteins involved in the regulation of organelle morphology, transport, and energetics

To determine possible host cell targets of SneRING, we expressed the FLAG-tagged protein in U2OS cells with and without Sne infection and performed a pull-down using FLAG-antibody followed by mass spectrometry analysis. We compared infected samples with and without ectopically expressed FLAG-tagged SneRING to determine significantly enriched proteins (Fig 5A). Highly enriched proteins (log2 fold change > 2) included mitochondrial proteins involved in organelle morphology (prohibitins, Mic60/IMMT), oxidative phosphorylation (OXPHOS) (subunits of ATPase and respiratory chain components), as well as transport (VDAC1 and 2). However, several ER proteins were also identified, and these included ER-microtubule anchoring protein CKAP4, BCAP31, a protein involved in ER-associated degradation (ERAD) and acting as an ER stress sensor that mediates communication with mitochondria (Namba, 2019), and ERLIN1 and 2, which are prohibitin-domain containing proteins involved in ERAD (Pearce *et al*, 2009) (Fig 5A, Table S1).

**Figure 5.**
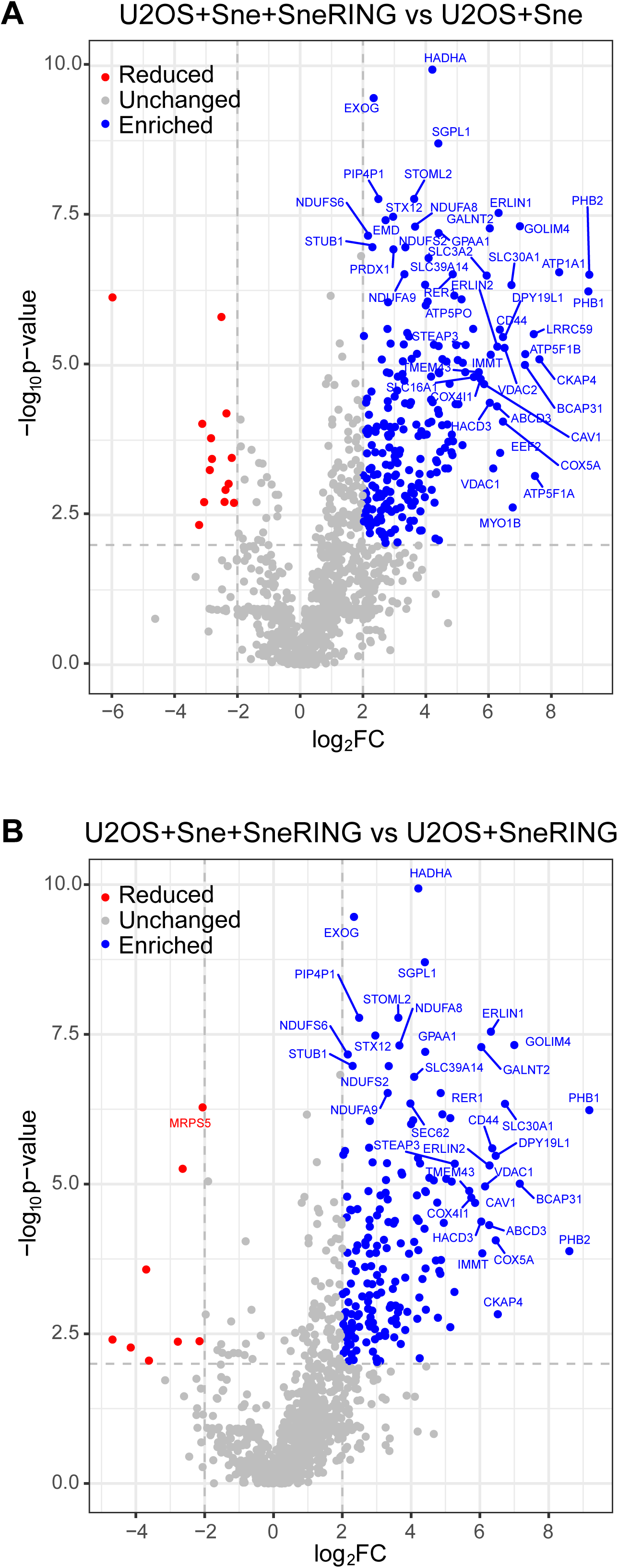
SneRING interacts with mitochondrial and ER proteins. **(A, B)** A human codon-adjusted FLAG-tagged version of the SneRING was overexpressed in U2OS cells using PEI MAX® transfection. 24 h later, cells were infected with Sne at an MOI 1. Non-transfected cells served as control. Immunoprecipitation was performed using FLAG-magnetic beads and the precipitated proteins were quantified by mass spectrometry. The graphs show identified proteins, with significance (-log_10_p-value) plotted against the log_2_ fold change (log_2_FC) of transfected/infected U2OS cells (U2OS+Sne+SneRING) relative to non-transfected/infected controls (U2OS+Sne) **(A)** or of transfected/infected U2OS cells (U2OS+Sne+SneRING) relative to transfected/non-infected controls (U2OS+SneRING) **(B)**. Enriched proteins are labeled in blue, reduced proteins are shown in red, and grey represents unchanged proteins. Only host cell proteins are shown.

Besides these cellular proteins, a number of Simkania-derived proteins were also strongly enriched in the SneRING-pulldowns (Fig S5, Table S1). These proteins might be bacterial SneRING targets or co- factors of SneRING. None of the top-enriched Sne proteins appears to be experimentally characterized, but several members of this set have informative sequence relationships. SNE_A02810, SNE_A02800, SNE_A02780, SNE_A18120, and SNE_A05090 are relatives of the Legionella major outer membrane protein mOMP, and SNE_08170 contains mOMP-associated POTRA domains. Other enriched proteins are related to components of bacterial secretion systems: A21850 is related to the SecDF secretion protein, SNE_A20110 is related to the T2SS secretin GspD, while SNE_A10070 is related to bacterial signal peptidases. Overall, the bacterial SneRING-associated proteins appear to be located at the bacterial surface.

When comparing the cellular SneRING-associated proteins of Sne-infected and non-infected cells, a very similar set of proteins was obtained. The top fourteen cellular infection-specific SneRING interactors were found also on the previous list. We again found mitochondrial and ER proteins among the hits. Interestingly, GOLIM4, a protein localized to Golgi apparatus and playing a role in endosome to Golgi protein transport (Natarajan & Linstedt, 2004), was also found as one of the most highly enriched proteins in both lists (Fig 5B, Table S1). Taken together, these results show clustering of putative SneRING targets in groups of ER and mitochondrial proteins functioning in quality control, bioenergetics, and morphology. The localization of these targets is in agreement with the possible localization of SneRING that we previously determined (Fig 4). The observation that most of the prominent SneRING interactors are only present in the infected samples suggests that SneRING needs additional bacterial factors for its function, or that the infection-induced remodeling of cellular compartments fosters SneRING interactions.

## Discussion

After invading the host cell, intracellular bacteria reside either directly within the cytosol or inside a vacuole. In both cases, they are targeted by the innate immune system of the host cell to induce their clearance and block their propagation. To survive and replicate, intracellular pathogens must modify and/or evade the host defense system. An important part of intracellular defenses is the ubiquitin system.

Ubiquitin system has been thought to be exclusive to eukaryotes; however, recent reports about bacterial ubiquitination systems that serve as anti-viral defense show that ubiquitination might be evolutionarily older than presumed (Chambers *et al*, 2024; Hör *et al*, 2024). In addition, various bacteria specifically counteract their ubiquitination and even hijack the system for their own advantage by secreting effector proteins (Tripathi-Giesgen *et al*, 2021). Thus, the ubiquitin system of the host cell represents a weapon for both the host cell and the bacteria. Bacterial DUBs can prevent bacterial ubiquitination, which can be a signal for xenophagy (Chai *et al*, 2019; Otten *et al*, 2021; Yamada *et al*, 2021). Furthermore, some bacteria deliver unique virulence factors that functionally mimic mammalian E3 ligases to exploit the host ubiquitin system. Ubiquitin E3 ligases of all three types have been described for various bacteria. These differ in their target in the host cell and their intracellular function (Ashida & Sasakawa, 2017).

Recent studies have shown that S. negevensis possesses a high number of DUBs from many different classes including some that have never before been seen in bacteria. This is in strong contrast to only one or two DUBs encoded by most other intracellular bacteria. Interestingly, some of the Simkania DUBs show cleavage specificity–SnOTU specifically cleaves linear M1 chains and VTD-type DUB preferably cleaves K6-linked ubiquitin conjugates–while others are less specific (Boll *et al*., 2023). These observations indicate that Sne encodes ubiquitin-modifying enzymes to manipulate and counteract the ubiquitin system of the host cell, but so far no Sne ubiquitin ligases have been described.

Bioinformatical analysis revealed several genes in Sne that contain a RING domain. In the case of SneRING, the RING as well as the C-terminal domain is shared with related proteins from other *Chlamydia*-like bacteria. Despite the overall similarity, there are notable differences in the Zn-binding residues of the RING finger, when comparing and aligning the sequences of the SneRING with the E3s from the *Chlamydia*-like bacteria. In addition to these putative E3s of related bacteria, also two established human RING fingers, structurally corresponding best to the SneRING model, were included in the alignment. In particular, PDB:3LRQ_A (from TRIM37) structurally matches best; however, it is more similar to the RING finger of the *Chlamydia*-like bacteria than of Sne in terms of Zn binding (Fig 1). In addition to SneRING, Sne possesses several more proteins with a RING domain (Fig S1), which remain to be characterized.

Though the RING finger of the SneRING appears to be rather atypical, we could demonstrate the ligase activity using autoubiquitination assays. SneRING does not unspecifically interact with E2 enzymes, because we detected the activity of the SneRING only in the presence of the E2 conjugating enzyme UbcH5b, and, to a lesser extent, in combination with the E2 UBE2T (Fig 2). However, the human genome encodes for around 40 E2 conjugating enzymes that are involved in transferring ubiquitin or ubiquitin-like proteins to the target (Stewart *et al*, 2016). We have tested only a limited number of E2 enzymes, so we cannot exclude that SneRING is active in the presence of other E2 enzymes besides UbcH5b and UBE2T.

UbcH5b plays an important role in various cellular processes, such as in the activation of NF-κB by the degradation of I_κ_bα (Gonen *et al*, 1999) and regulation of the level of p53 by ubiquitination and its subsequent degradation (Saville *et al*, 2004). It is involved in the removal of abnormal and regulatory proteins (Seufert & Jentsch, 1990) and is one of the most active E2 enzymes (Houben *et al*, 2004). In contrast, the E2 conjugating enzyme UBE2T regulates the levels of the receptor for activated protein kinase C (RACK1) by mediating its ubiquitination with K48-linked conjugates followed by its elimination. In this way, UBE2T induces hyperactivation of the Wnt/β-catenin signaling pathway (Yu *et al*, 2021). The Wnt/β-catenin signaling pathway is crucial in various physiological processes ranging from embryonic development to tissue homeostasis in adults (Liu *et al*, 2022). It is conceivable that SneRING interacts with these E2 ligases during infection, considering their ubiquitous presence in the cell. 3D structure prediction has helped identify S19 and E22 as crucial for the interaction with UbcH5b, which was confirmed by an *in vitro* autoubiquitination assay (Fig 1C and Fig 2B). This shows that the predicted RING domain of the SneRING is the interacting region with the E2 conjugating enzyme UbcH5b. The expression of the catalytically inactive form of SneRING, however, did not influence infection.

SneRING produces K6-, K11-, K48- and K63-linked ubiquitin conjugates, which implies that its host cell targets are modified similarly (Fig 3 and Fig S2). The preference to catalyze K6 and K48 linkages was also observed for the HECT-like ubiquitin ligase NleL of Escherichia coli O157:H7. (Lin *et al*, 2011). This strain from the group of enterohemorrhagic E. coli (EHEC) causes hemorrhagic colitis and diarrhea in humans (Lim *et al*, 2010). The bacterial infection is stimulated by the bacterial HECT-like ligase NleL, which mediates the ubiquitination and inactivation of the host c-Jun NH2-terminal kinases (JNKs), promoting the production of unique attaching and effacing (A/E) lesions (Sheng *et al*, 2017) and thus a close adherence between the bacterium and the membrane of the epithelial cell (Nataro & Kaper, 1998). NleL shows *in vitro* ubiquitinating activity when incubated with the E2 conjugating enzyme UbcH5b (Lin *et al*., 2011), similar to SneRING.

Although not obligate intracellular, Salmonella enterica subsp. enterica serotype Typhimurium replicates within a bacteria-containing vacuole after invading the host cell (Steele-Mortimer, 2008), similar to Sne. SopA effector of S. Typhimurium is a HECT-like E3 ubiquitin ligase and preferentially reacts with the host E2 ubiquitin-conjugating enzymes UbcH5a, UbcH5c, and UbcH7. It has been proposed that SopA ubiquitinates bacterial and host cell proteins involved in Salmonella-induced intestinal inflammation (Zhang *et al*, 2006). The ligase selectively mediates K11- and K48-linked polyubiquitination of the human RING-ligases TRIM56 and TRIM65, leading to their proteasomal degradation during infection. Blocking these ligases suppresses the expression of the nucleic acid-sensing receptor STING and interferon-γ (Fiskin *et al*, 2017; Tripathi-Giesgen *et al*., 2021). S.

Typhimurium virulence factor SspH2 has been shown to autoubiquitinate *in vitro* when combined with the E2 enzyme UbcH5b and is thus assumed to be an E3 ubiquitin ligase, too, similar to HECT type E3 ligases (Quezada *et al*, 2009). *in vitro* experiments revealed that SspH2 mediates K48-linked ubiquitination, suggesting that it targets proteins for degradation by the proteasome (Levin *et al*, 2010). A putative homolog of SspH2, IpaH3 from Shigella, also mediates the synthesis of K48-linked ubiquitin chains and it only shows activity in the presence of E2 conjugating enzymes of the UbcH5 type (Zhu *et al*, 2008). Ultimately, all the observations made in this study and parallels with known E3 ligases of other bacteria support the identification of SneRING as an E3 ubiquitin ligase with a role in infection. Since genetic modification of Sne has so far not been achieved, we are currently unable to confirm to which extent bacterial infectivity and life cycle depend on the activity of SneRING.

The variety of Sne DUBs, with 12 identified candidates of which 8 are active, indicates that ubiquitin-modifying enzymes are important for Sne infection. As with other intracellular bacteria, Sne replication niche SnCV can be a direct target for ubiquitination. This could lead to its complete or partial degradation by autophagy (Hermanns & Hofmann, 2019). Sne DUBs might play an important role in preventing autophagy by removing ubiquitin from the SnCV surface. The role of SneRING, on the other hand, can be to modulate protein stability or activity through ubiquitination. While some Sne DUBs do not show cleavage specificity, others are only active against certain ubiquitin chain types. Interestingly, the two Sne-encoded DUBs SnJos1 and SnJos2 efficiently cleave K6-, K11-, K48- and K63-linked ubiquitin chains, with another enzyme selectively cleaving K6-linked chains (Boll *et al*., 2023). One possibility is that SneRING generates these specific ubiquitin conjugates in the early stages of infection for them to be removed at later stages by Sne DUBs to support bacterial development and/or release. However, whether the targets of both SneRING and Sne DUBs are bacterial or host cell proteins remains yet to be determined.

SnCV has a unique tubular morphology and is in close contact with ER. Its development depends on retrograde vesicular transport (Herweg *et al*., 2016) and it is presumed that the majority of lipids for growth of this complex membranous compartment are derived from ER. Sne is also capable of suppressing ER stress response (Mehlitz *et al*., 2014). On the other hand, mitochondria intertwine with SnCV only in the early stages of infection, whereas at the later time point, we observe fragmentation and loss of mitochondrial mass. Endogenous SneRING is found in the ER/SnCV membrane (Herweg *et al*., 2016), whereas the overexpressed SneRING localizes to cytosol, ER, and mitochondria (Fig 4 and Fig S4), including its surprising association with the inner mitochondrial membrane (Fig 4D). This, together with the identification of possible targets and interacting factors, might offer a clue about the function of this enzyme.

Mitochondrial depolarization leads to the formation of K6-, K11-, K48-, and K63-chain linkages on the surface of these organelles. Parkin, a mitochondrial protein with E3 ubiquitin ligase activity and the potential to form these same linkages is involved in mitochondrial ubiquitination, which promotes mitophagy (Harper *et al*, 2018). Recently, Parkin has been shown to ubiquitinate proteins not only on the surface of mitochondria but also in the inner membrane. Parkin directly binds to prohibitin 2 (PHB2) through its RING1 domain and promotes K11- and K33-linked ubiquitination of PHB2 (Sun *et al*, 2022), which enhances the interaction between PHB2 and the central protein in autophagy, LC3 (Wei *et al*, 2017). SneRING not only generates the same types of ubiquitin chains in an *in vitro* reaction as Parkin (Fig. 3), but one of the highly enriched host cell proteins in the SneRING pulldown fractions is PHB2 (Fig 5, Table S1). SneRING might ubiquitinate PHB2 to induce mitophagy; however, this process would depend on the presence of SnCV and other, so far unidentified bacterial factors, since we did not observe mitochondrial fragmentation in non-infected cells expressing SneRING (Fig S4). Alternatively, SneRING/prohibitin interaction might impact mitochondrial integrity and respiration, in which prohibitins play an important role (Merkwirth & Langer, 2009). This could also explain another highly enriched putative SneRING interactor, BCAP31, a protein present in ER-mitochondria contact sites, which has been reported to regulate mitochondrial respiration by controlling the mitochondrial import of respiratory complex I subunits (Namba, 2019). Interestingly, ERLIN1 and ERLIN2, other highly enriched hits (Fig 5, Table S1), also contain a prohibitin domain. These proteins localize to ER and are involved in ERAD (Pearce *et al*., 2009), which might be a potential target of modification by SneRING considering that bacteria can suppress ER stress response (Mehlitz *et al*., 2014), a process counteracted by the ERAD upregulation (Hwang & Qi, 2018). Finally, possible SneRING ubiquitination of CKAP4, a protein involved in the attachment of ER to microtubules (Vedrenne *et al*, 2005), might be required during the growth and organization of the SnCV. CKAP4 has recently been reported to also play a role in ER-phagy, a process triggered by ER stressors, such as tunicamycin (Li *et al*, 2024). Its targeting by SneRING might be one of the ways for bacteria to interfere with ER stress signaling (Mehlitz *et al*., 2014). Though interesting, these theories must be confirmed by analyzing if any of these proteins are ubiquitinated upon SneRING expression or during infection. In addition, deletion of SneRING from bacteria could give further information about the function and importance of this protein.

To survive and replicate within their host, intracellular pathogens have evolved various strategies and mechanisms to counteract the host cell immune system. By secreting effector proteins, pathogens manipulate the host cell to provide better conditions for their survival and propagation. Effector proteins also include ubiquitin-modifying enzymes such as deubiquitinases and ubiquitin E3 ligases. This study describes for the first time a Sne effector protein SneRING that functions as ubiquitin E3 ligase with a possible role in remodeling host cell mitochondria and ER and controlling mitophagy and ER stress response to promote the development of SnCV and accommodate its unique replicative niche.

## Materials and Methods

### Sequence analysis

Sequence alignments were generated using the MAFFT package (Katoh & Standley, 2013). Generalized profiles were derived from multiple alignments using pftools (Bucher *et al*., 1996) and searched against the Uniprot database (https://www.uniprot.org) and the NCBI microbial genome reference sequence database (https://www.ncbi.nlm.nih.gov/genome/microbes) using pfsearchV3 (Schuepbach *et al*, 2013). Structure predictions were run using a local installation of Alphafold 2.3 (Jumper *et al*., 2021). Structure comparisons were performed using the DALI software (Holm, 2022). For modeling the missing Zn^2+^ ions into Alphafold structures, a superposition with the best-scoring Zn^2+^-containing DALI hit was used to copy the zinc ions into the model. Subsequently, an energy minimization was performed using YASARA (Krieger & Vriend, 2014) to improve the sidechain positioning of the Zn^2+^ ligands.

### Cloning and mutagenesis

The SneRING coding region for protein purification was obtained by amplification of S. negevensis genomic DNA (Leibniz Institute DSMZ, German Collection of Microorganisms and Cell Cultures, DSM No. 27360) using the following primers: forward, 5’- AAGTTCTGTTTCAGGGCCCGGAAAGGGTAAATCCTAATCAAGTTCTA – 3’, and reverse, 5’-ATGGTCTAGAAAGCTTTATGAAAAAATTCTACGAGAAATATCTTCC – 3’. Amplification was performed using the Phusion^TM^ High Fidelity DNA Polymerase Kit (Thermo Fisher Scientific, Massachusetts, USA), and the PCR product was inserted into the pOPIN-K vector (Berrow *et al*, 2007) by restriction cloning according to the manufacturer’s protocol. Point mutations were generated using the QuikChange Lightning kit (Agilent Technologies, California, USA). To introduce the first mutation, the following primers were used: forward, 5’ – GCATGGAATTGTCGCAGGTGCTGGGAACCAATTCA – 3’, and reverse, 5’ – TGAATTGGTTCCCAGCACCTGCGACAATTCCATGC – 3’. The second mutation was added by using the following primers: forward, 5’ – AATTGTCGCAGGTGCTGGAGACCAATTCAGGATGGTCC – 3’, and reverse, 5’ – GGACCATCCTGAATTGGTCTCCAGCACCTGCGACAATT – 3’. SneRING coding region for transfection experiments was synthesized by Thermo GeneArt Gene synthesis (Thermo Fisher Scientific, Massachusetts, USA) and cloned into the pCDNA3 vector by restriction digestion according to the manufacturer’s protocol.

### Protein expression and purification

Wildtype SneRING was expressed from the pOPIN-K vector containing an N-terminal 6His-GST-tag in E. coli Rosetta (DE3) pLysS. The bacteria culture was grown in LB medium at 37 °C until reaching an OD_600_ of 0.8. Afterward, the culture was precooled at 18 °C before SneRING expression was induced with 500 µM ZnCl_2_ and 0.1 mM isopropyl β-d-1-thiogalactopyranoside (IPTG) for 16 h when bacteria were collected by centrifugation at 5,000 x g for 15 min. The pellets were frozen at -80 °C, then thawed on ice and resuspended in binding buffer (300 mM NaCl, 20 mM Tris pH 7.5, 20 mM imidazole, 2 mM β-mercaptoethanol), containing DNase and lysozyme. Cells were sonicated for 10 min with 10 s pulses at 50 W and the lysate was clarified by centrifugation for 1 h at 50,000 x g and 4 °C. The supernatant was applied to a HisTrap FF column (Cytiva, Massachusetts, USA), and affinity purification was performed according to the manufactureŕs protocol. The subsequent incubation of the fractions with 3C protease during dialysis in binding buffer overnight resulted in the 6His-GST tag removal, which was removed, together with the His-tagged 3C protease in a second affinity purification run. Size exclusion chromatography (HiLoad 16/600 Superdex 75 pg (Cytiva, Massachusetts, USA)) was performed in a buffer containing 20 mM Tris pH 7.5, 150 mM NaCl, and 2 mM dithiothreitol (DTT). After concentration using VIVASPIN 20 Columns (Sartorius, Göttingen, Germany), the purified protein was frozen in liquid nitrogen and stored at -80 °C.

The expression of SneRING S19R_E22R was induced as already described for the wildtype SneRING. After harvesting the cells, the pellet was resuspended in lysis buffer (40 mM Tris pH 8.2, 500 mM NaCl, 10 mM β-mercaptoethanol, 10 mM imidazole, containing 1xComplete^TM^ protease inhibitor cocktail (Roche Holding, Basel, Switzerland), DNase and lysozyme). Cells were lysed by sonication, the cleared lysate was applied to Ni-NTA agarose (Qiagen, Hilden Germany), and affinity purification was performed according to the manufactureŕs protocol. Mutated SneRING was eluted using 40 mM Tris pH 7.5, 500 mM NaCl, 10 mM β-Mercaptoethanol, and 250 mM imidazole. The removal of the 6His-GST tag was performed similarly to the wildtype SneRING and using the Ni-NTA agarose (Qiagen, Hilden Germany) in a buffer containing 20 mM Tris pH7.5 and 300 mM NaCl. After concentration on Amicon Ultra-15, PLGC Ultracel-PL membrane (Sigma/Merck, Darmstadt, Germany), the protein was frozen in liquid nitrogen and stored at -80 °C.

The two Sne DUBs SnVTD and SnOTU used in the study were purified as described before (Boll *et al*., 2023). The plasmids pOPINB-OTUB1* (Addgene plasmid #65441; http://n2t.net/addgene:65441; RRID:Addgene_65441), pOPINB-AMSH* (Addgene plasmid #66712; http://n2t.net/addgene:66712; RRID:Addgene_66712) (Michel *et al*, 2015), pOPINK-Cezanne (OTU, aa 53-446), (Addgene plasmid #61581; http://n2t.net/addgene:61581 ; RRID:Addgene_61581) (Mevissen *et al*, 2013) and pOPINS-USP21 (USP, aa 196-565) (Addgene plasmid #61585; http://n2t.net/addgene:61585; RRID:Addgene_61585) (Ye *et al*., 2011) were kind gifts from David Komander (WEHI, Melbourne, Australia). Purification of these DUBs was performed as already described for the SneRING. Protein concentration was determined by measuring the absorption at 280 nm (A_280_) and using the SneRING extinction coefficient calculated from its sequence.

### *in vitro* autoubiquitination assay

10 µL of the 10x ligation buffer (800 mM Tris pH 7.5, 200 mM MgCl_2_, 12 mM DTT) was mixed with 10 µM purified recombinant SneRING, 100 nM recombinant E1 ubiquitin-activating enzyme and 2.3 µM recombinant E2 ubiquitin-conjugating enzyme, to which 50 µM recombinant ubiquitin and 10 mM ATP were added. ddH_2_O was added to a final volume of 100 µL. The reactions were incubated at 37 °C for different time points (0 min, 30 min, 60 min, 120 min, 180 min, and 300 min) and stopped by adding Laemmli sample buffer (62.5 mM Tris pH 6.8, 2% SDS, 10% glycerol, 5% β-mercaptoethanol, and 0.002% Bromophenol Blue) and analyzed by sodium dodecyl sulfate-polyacrylamide gel electrophoresis (SDS-PAGE) and western blot.

The purified proteins of the E2 conjugating enzymes UbcH5b, UbE2T, UbCH7, Ube2A, Ube2B and Ube2W were a kind gift from David Komander (WEHI, Melbourne, Australia).

### UbiCRest assay

The recombinant DUBs were first diluted in a buffer containing 150 mM NaCl, 20 mM Tris pH 7.5, and 10 mM DTT. The 5 h *in vitro* autoubiquitination assay reaction was treated for 15 min at RT with 2 mU apyrase to remove the remaining ATP. For the assay, the DUBs were added at 1.5 µM concentration except OTUB1, for which 3 µM concentration was used. During the double UbiCrest assay, the DUBs were added at 1.5 µM or 0.75 µM concentrations. The mixture was incubated for 1 h at RT and the reaction was stopped by adding Laemmli sample.

### Immunoblotting

The lysates for western blot analysis were prepared with Laemmli sample buffer and denatured at 95 °C for 5 min. After SDS-PAGE, proteins were transferred onto a PVDF membrane. The membrane was blocked with 5% milk in Tris-based saline (TBS) and incubated in the primary antibody, followed by the incubation with an HRP-coupled secondary antibody (1:3000) in a blocking solution. The signal was detected using SuperSignal™ West Femto Maximum Sensitivity Substrate (Thermo Fisher Scientific, Massachusetts, USA) and a Chemo Cam Imager (Intas). In the case of an affimer, the membrane was incubated overnight at 4 °C in the affimer dilution in TBS. The next day, the membrane was washed with TBS and incubated with a 6xHis primary antibody, followed by the HRP-coupled secondary antibody as already described.

### Cell culture

HeLa (ATCC® CCL-2.1^TM^), HeLa229 KDEL-dsRed, Hela 229 Mito-GFP, and THP-1 (ATCC® TIB-202^TM^) cells were grown in Roswell Park Memorial Institute (RPMI)1640 medium (Thermo Fisher Scientific, Massachusetts, USA), Hek293T cells (ATCC® CRL-3216^TM^) were grown in Dulbecco’s modified Eagle’s medium (DMEM) medium (Thermo Fisher Scientific, Massachusetts, USA) and U2OS cells (ATCC® HTB-9.6^TM^) were grown in DMEM/F12 medium (Thermo Fisher Scientific, Massachusetts, USA). All media were supplemented with 10% v/v heat-inactivated (56 °C at 30 min) fetal calf serum (FCS) (Sigma/Merck, Darmstadt, Germany). Growth took place at 37 °C and 5% CO_2_. To differentiate THP-1 cells into macrophages, 5x 10^5^ cells/well were seeded into 6-well plates and incubated with 20 ng/mL phorbol 12-myristate 13-acetate (PMA) (Sigma/Merck, Darmstadt, Germany) in RPMI1640 medium containing 10% v/v heat-inactivated FCS for 24 h. The medium was exchanged for fresh RPMI1640 medium containing 10% v/v heat-inactivated FCS and the cells were incubated for a further 48 h.

Primary human macrophages were derived from peripheral blood mononuclear cells (PBMCs) isolated from leukoreduction system (LRS) cones, using the SepMate™-50 system (#85450, STEMCELL Technologies, Cologne, Germany) and Ficoll-Paque (#GE17-1440-03, Sigma/Merck, Darmstadt, Germany) gradient according to manufacturer’s instructions. Red blood cells were removed by hypo-osmotic shock: PBMC pellets were mixed and incubated with 16 mL sterile water for 30 s, then immediately 4 mL 5 x DPBS were added. Monocytes were purified from the PBMC fraction using the EasySep™ Human CD14 Positive Selection Kit II (# 17858, STEMCELL Technologies, Cologne, Germany) according to the manufacturer’s instructions and seeded in 6-well cell culture plates (1.5 x 106 cells/well) in medium supplemented with 50 ng/mL M-CSF (#78057, STEMCELL Technologies, Cologne, Germany). Medium exchanges, with fresh M-CSF supplementation, were done on days 1 and 4. On day 7, M2”-like polarization, was induced by replacing M-CSF-containing medium with medium containing 100 ng/mL IL-4 (#78045, STEMCELL Technologies, Cologne, Germany) and further incubation for 48 h. All cultivation steps were done in RPMI1640 (Thermo Fisher Scientific, Massachusetts, USA) supplemented with 10% (v/v) heat-inactivated FCS (Sigma/Merck Darmstadt, Germany). All Sne infection experiments were performed on day 9, without cytokine supplementation.

Acanthamoeba castellanii (ATCC® 30010D^TM^) were cultured in axenic medium (56 mM glucose, 2.6 mM KH2PO4, 1.3 mM Na2HPO4x7H2O, 0.5% proteose peptone 2, 0.5% thiotone E peptone, 0.5% yeast extract) in cell culture flasks at 30 °C.

### Bacteria and infection

A. E. coli DH5α and Rosetta (DE3) pLysS were grown in an LB medium containing the respective antibiotics for selection (1% bacto-tryptone, 0.5% yeast extract, 1% NaCl, 1% agar).

To prepare S. negevensis (ATCC® VR-1471^TM^), U2OS cells were grown to 60% confluency, and infected at a multiplicity of infection (MOI) 1 in RPMI1640 medium without HEPES containing 5% v/v heat-inactivated FCS (infection medium). The infected cells were incubated at 35 °C and 5% CO_2_ for 6 h, when the medium was exchanged for fresh infection medium. After 3 days of growth under the same conditions, the cells were detached using a cell scrapper and lysed by vortexing the cells with 2-5 mm glass beads (Carl Roth, Karlsruhe, Germany). Intact cells and larger cell debris were removed by low-speed centrifugation at 600 x g for 10 min and the resulting supernatant was centrifuged at 20,000 x g for 30 min to pellet the released bacteria. The pellet was resuspended in 5 mL SPG buffer (250 mM sucrose, 4 mM monopotassium phosphate, 10 mM disodium phosphate, and 5 mM glutamate pH 7.4), followed by centrifugation (20,000 x g, 30 min), with the resulting pellet resuspended in 2.5 mL SPG buffer. The bacteria were homogenized using a syringe with one G20 and one G18 needle, aliquoted, and stored at -80 °C. All the experiments with S. negevensis were performed in a laboratory with biosafety level 2, which is registered with the Lower Franconia Government under code 55.1-8791.1.30.

For infection experiments, 5 x 10^5^ human cells/well were seeded into 6 well plates and at 60% confluency infected with S. negevensis at MOI 1 in the infection medium, at 35 °C and 5% CO_2_. 6 h later, the medium was exchanged for fresh infection medium. The infection proceeded for the indicated time points. When infection took longer than 3 days, the medium was exchanged for fresh infection medium after every 3 days.

A. castellanii were seeded into cell culture flasks and infected with S. negevensis in an axenic medium. Infected amoebae were incubated at 30 °C for 4 days.

### Transfection

Hek293T cells were transfected using the Calcium-phosphate transfection method as described previously (Wagner *et al*, 2019). For 6-well plates, 4 µg of the respective plasmid were used; for transfecting cells in 15 cm dishes, we used 50 µg of plasmid DNA. U2OS and HeLa229 cells were transfected using the commercially available transfection reagents jetOptimus® (Polyplus®, Illkirch-Graffenstaden, France) or PEI MAX® (Polysciences, Inc., Warrington, United Kingdom) according to the manufacturer’s protocol. Depending on the experiment, cells were infected with S. negevensis at MOI 1, 24 h post-transfection.

### Immunofluorescence and Microscopy

U2OS and HeLa cell lines were seeded at a density of 8 x 10^4^ cells/well into 12 well plates, while Hek293T cells were seeded at a density of 5 x 10^5^ cells/well into 6 well plates on 15 mm glass coverslips (Paul Marienfeld, Lauda-Königshofen, Germany). Cells were transfected and/or infected as indicated in the corresponding experiments. The cells were fixed at respective time points using 4% paraformaldehyde (PFA) in PBS for 1 h at RT. For S. negevensis infection experiments, the cells were permeabilized with 0.2% Triton X-100 in PBS for 45 min at RT while shaking, washed with PBS, and blocked in 2% goat serum in PBS for 1 h, followed by staining with a SnGroEL primary antibody in the blocking solution overnight at 4 °C. The following day, samples were washed and incubated with the secondary antibody (1:1600) and DAPI (1:3000) for 1 h in 2% goat serum in PBS at RT. After washing with PBS and ddH_2_O, coverslips were mounted using Mowiol 4-88 (Carl Roth GmbH + Co. KG, Karlsruhe, Germany).

For experiments that did not include infection, after fixing, cells were simultaneously permeabilized and blocked for 30 min at RT in blocking solution (10% BSA, 0.2% Triton X-100, in PBS), washed with PBS and stained with the primary antibody diluted in antibody dilution solution (3% BSA in 0.05% Tween in PBS) for 1 h at RT. After washing with PBS and blocking for 10 min in blocking buffer at RT, samples were washed again and incubated with DAPI (1:3000) and the secondary antibodies (1:100) for 1 h at RT. The coverslips were then washed with PBS and ddH_2_O and mounted using Mowiol 4-88 (Carl Roth GmbH + Co. KG, Karlsruhe, Germany). Microscopy was performed using LEICA SP5 confocal laser scanning microscope.

### RNA isolation, cDNA synthesis, and RT-qPCR

Infected macrophages were collected on day 2, 4 and 6 pi. The cells were washed with DPBS at the desired time point and 100 µL of fresh medium and 500 µL RNAprotect Cell Reagent (Qiagen, Hilden, Germany) were added per well. Detached cells were collected in RNase-free tubes and stored at 4 °C for a maximum of 1 week. After centrifugation at 6200 x g for 5 min at 4 °C, 600 µL RLT Buffer supplemented with β-mercaptoethanol was added to each pelleted sample, and RNA was purified using the RNeasy Mini kit (Qiagen, Hilden, Germany) according to manufacturer’s instructions. For other human cell lines, after collecting them on day 2 and 4 pi and washing with DPBS at the desired time point, 600 µL RLT Buffer supplemented with β-mercaptoethanol was added to each sample and RNA was purified using the RNeasy Mini (Qiagen, Hilden, Germany) according to manufacturer’s instructions. DNase treatment was performed using the TURBO DNA-free™ Kit (Invitrogen, California, USA). 100 ng pure RNA was used for cDNA generation, using the Revert Aid First Strand cDNA Synthesis Kit (Thermo Fisher Scientific, Massachusetts, USA). RT-qPCR was performed on a StepOnePlus™ instrument (Life Technologies, California, USA), using the GreenMasterMix (2X) High ROX™ (Gennaxon, Ulm, Germany). Data analysis was performed using the StepOne Software (v2.3) and Microsoft Excel, with a modified 2 ^-ΔΔCt^) method (Livak & Schmittgen, 2001). ΔCt values were calculated relative to the host reference gene YWHAZ (human) or the 5S RNA (for amoeba). Fold changes (2^-ΔΔCt^) of gene expression for each tested Sne gene, were calculated relative to the non-infected controls. The following primers were used: forward, 5’ – ATACTGCGCAGAAAGCAC – 3’, and reverse, 5’ – ACCCCAGTACTAACACCG – 3’ (5S RNA reference gene for amoeba); forward, 5’ – GCAGAGTCGAAGTTTCAAGGG – 3’, and reverse, 5’ – TCGCTTGCCTGTTGAGGATT – 3’ (SneRING); forward, 5’ – TTCCATCACCGGCAACATCA – 3’, and reverse, 5’ – GCAAAAGAGATCGCGCTGAA – 3’ (GroEL C (SNE_A22590)); forward, 5’ – CACCTGATCCCATCCCGAAC – 3’, and reverse, 5’ – GCGACCTACTCTCCCGTACT – 3’ (Sne 5S RNA) (Boll *et al*., 2023); forward, 5’ – ACTTTTGGTACATTGTGGCTTCAA – 3’, and reverse, 5’ – CCGCCAGGACAAACCAGTAT – 3’(YWHAZ) (Vandesompele *et al*, 2002).

### Mitochondrial isolation and fractionation

Mitochondria were isolated from cells transfected for 48 h as described previously (Humphries *et al*, 2005). Cellular fractionation and determining of submitochondrial localization by swelling and PK treatment were performed as already described (Kozjak-Pavlovic *et al*, 2011; Kozjak *et al*, 2003). Carbonate extraction for determining protein membrane extractability was performed as previously described (Milenkovic *et al*, 2004). In brief, isolated mitochondria from transfected cells were either sonicated in 10 mM Tris pH 7.6 and 500 mM NaCl or treated with 100 mM Na_2_CO_3_ at pH 10.8 or 11.5 for 20 min on ice. After centrifugation at 100.000 g, pellets were resuspended in Laemmli sample buffer, while supernatant and non-centrifuged control samples were precipitated using trichloroacetic acid (TCA).

### Antibodies, chemicals, and positive controls

K48- and K63-linked ubiquitin chains were components of the Di-ubiquitin Explorer Panel purchased from UbiQ (Amsterdam, Netherlands, Cat. No. UbiQ-L01). M1-linked ubiquitin chains (kindly provided by S. Meyer and A. Buchberger) were produced by incubating the following reaction mixture at 37 °C for two hours: 6 µg bovine ubiquitin, 200 ng yeast Uba1, 600 ng human UbcH5c, 4 µg HOIP RBR-LDD (Smit *et al*, 2012) in a buffer containing 50 µL 20 mM Tris/HCl pH 7.5, 5 mM MgCl_2_, 1 mM DTT, 4 mM ATP.

The primary antibodies against ubiquitin (Cat. No. 05-9449), FLAG-tag (Cat. No. F7425), CPOX (Cat. No. HPA015736), K63-linked ubiquitin chains (Cat. No. 05-1308), K48-linked ubiquitin chains (Cat. No. 05-1307) and M1-linked ubiquitin chains (Cat. No. MABS451, kindly provided by A. Buchberger) were purchased from Sigma/Merck (Darmstadt, Germany). The anti-ubiquitin Lys6 specific Affimer® (Cat. No. MABS1918, kindly provided by M. Eilers) was purchased from Sigma/Merck (Darmstadt, Germany). The primary antibodies against Tubulin (Cat. No. ab18251) and Mitofilin (Cat. No. ab48139) were purchased from Abcam (Cambridge, United Kingdom). The primary antibodies against Calnexin (Cat. No. 2433, Cat. No. sc-23954) were purchased from Cell Signaling (Massachusetts, USA) and Santa Cruz Biotechnology (Texas, USA). The antibody against GAPDH (Cat. No. sc-47724) was purchased from Santa Cruz Biotechnology (Texas, USA). The anti-HSP60 antibody (Cat. No. ADI-SPA- 806) was purchased from Enzo Life Sciences (New York, USA). The primary antibody against TIM44 (Cat. No. 612582) was purchased from BD Transduction Laboratories (California, USA). The antibody against 6xHis (Cat. No. GTX18184) was purchased from Biozol (Eching, Germany). The antibody against K11-linked ubiquitin chains (Cat. No. NBP3-05681, kindly provided by A. Buchberger,) was purchased from Novus Biologicals (Littleton, USA). The antibody against Sam50 was custom-generated against the full-length protein (Gramsch laboratories, Schwabhausen, Germany). The antibody against rat Tom70 (crossreactive with human Tom70) was a gift from K. Mihara. The antibody against SnGroEL was produced as described in a previous study (Mehlitz *et al*., 2014). Secondary antibodies anti-rabbit-Alexa555, Cat. No. A-21428, anti-mouse-Alexa488 Cat. No. A-11001, and anti-mouse-Alexa555, Cat. No. A-21422 were purchased from Thermo Fisher Scientific (Massachusetts, USA). Anti-rabbit-Atto647N secondary antibody, Cat No. 40839, was purchased from Sigma/Merck (Darmstadt, Germany). Anti-mouse-Cy2 secondary antibody, Cat. No. 115-225-146, as well as the HRP-coupled secondary antibodies for western blot, were purchased from Biozol (Eching, Germany).

### Immunoprecipitation

For immunoprecipitation (IP) assays, U2OS cells were seeded at a density of 1.2 x 10^6^ cells/well into 10 cm dishes. The next day, the cells were transiently transfected with a plasmid carrying a FLAG-tagged humanized version of the SneRING gene and 24 h later infected with Sne at an MOI 1. Non-transfected and non-infected cells served as controls. On day 2 pi, the cells were washed once with PBS, harvested in lysis buffer (20 mM Tris pH 7.5, 150 mM NaCl, 0.1 % NP-40, proteinase inhibitor (Roche: cOmplet, EDTA-free Protease Inhibitor Cocktail), 1 mM PMSF, 5 mM Iodoacetamide)), incubated for 30 min at 4 °C and sonicated for 15 min using an ultrasonic water bath. After removing the cell debris by centrifugation (15 min, 14000 x g, 4 °C), the supernatant was added to equilibrated Anti-FLAG® M2 magnetic beads (Sigma/Merck, Darmstadt, Germany) and incubated on a rotary shaker overnight at 4 °C. The beads were washed three times for 5 min at 4 °C with washing buffer (20 mM Tris pH 7.5, 150 mM NaCl, 0.1% NP-40) and once with a buffer containing 20 mM Tris pH 7.5, 150 mM NaCl. For mass spectrometry analysis, bound proteins were eluted with 200 µg/mL FLAG peptide (Sigma/Merck, Darmstadt, Germany) in elution buffer (6 M urea, 2 M thiourea).

### Mass spectrometry analysis

For in-solution digest, the samples were reduced by adding 1 µL of 1 M DTT for 1 h at RT, alkylated with 1 µL of 400 mM chloroacetamide (CAA) for 45 min at RT in darkness, digested with 1 µL of the Lysyl endopeptidase (Lys-C, 0.5 µg/µL) for 3 h at RT, treated with 270 µL of 50 mM triethylammonium bicarbonate buffer and finally digested with 1 µL of trypsin enzyme (1 µg/µL) overnight at RT. The next day, samples were acidified by adding 1 µL of 100% formic acid. After in-solution digestion, peptides were bound to styrene-divinylbenzene reverse-phase stage tips (SDB-RP). One stage tip was used per sample. The stage tips were equilibrated by adding 20 µL of 100% methanol and centrifuging for 1 min at 700 x g. Afterward, 20 µL of buffer B (0.1% formic acid in 80% acetonitrile) was added and the stage tips were again centrifuged for 1 min at 700 x g. In the next step, 20 µL of buffer A (0.1 % formic acid in water) was added to each stage tip, the samples were centrifuged for 1 min at 700 x g and an additional 20 µL of buffer A was added. Acidified samples were spun down for 5 min at 17,000 x g, the supernatant was loaded onto the SDB discs of the equilibrated stage tips, and they were again centrifuged for 5 min at 700 x g. The stage tips were washed buffers A and B and completely dried with a syringe.

After binding peptides to SDB-RP Stage tips, they were analyzed by liquid chromatography-mass spectrometry/mass spectrometry (LC-MS/MS). LC-MS/MS analysis was performed by the Proteomics Facility of the CECAD, University of Cologne, as described previously (Hermanns *et al*, 2018).

## Acknowledgments

We thank Prof. Dr. David Komander for the University of Melbourne for providing a panel of deubiquitinating and E2 ubiquitin-conjugating enzymes. We thank Prof. Dr. Alexander Buchberger and Prof. Dr. Martin Eilers from the University of Würzburg and Prof. Dr. Katsuyoshi Mihara from the Kyushu University for antibodies. We also thank Susanne Meyer for the University of Würzburg for the production of M1-linked ubiquitin chains. This study was supported by the Deutsche Forschungsgemeinschaft (DFG, German Research Society) through a GRK2243/1+2 grant to VK-P.

## Ethics statement

Primary cells used in this study originate from leukoreduction system cones made available by the Institut für Klinische Transfusionsmedizin und Hämotherapie of the University of Würzburg, Germany for research purposes and are not subject to approval by the Ethics Committee of the University of Würzburg.

