## Supplemental Figure Legends for "Identification and characterization of a ubiquitin E3 RING ligase of the *Chlamydia*-like bacterium *Simkania negevensis*"

**Supplementary Figure and Table legend:**

**Supplementary Figure 1.** **Sne_A08700 and Sne_A19470 are two other RING-like E3 ligase candidates from *S. negevensis*. (A)** Alphafold model of Sne_A08700 shows an N-terminal RING-like domain (blue) with a large insertion after the third Zn-coordinating residue (grey). The Zn_1_ and Zn_2_ ions coordinated by the RING-like domain are shown in red and magenta, respectively. **(B)** Multiple alignment of the RING domains of Sne_A08700, Sne_A08690, and some bacterial relatives (Chlr-KDK64: *Chlamydiia* bacterium, Uniprot: A0A960RSF6; Ver-JSS10: *Verrucomicrobiota* bacterium, Uniprot: A0A9E0XGC6; Neptunochl: Candidatus *Neptunochlamydia* sp., RefSeq: WP_316358199) and the sequence of the best DALI hit (pdb:5DKA). Residues invariant or conserved in at least 50% of the sequences are shown on black and grey background, respectively. Residues involved in the coordination of Zn_1_ and Zn_2_ are highlighted in red and magenta, respectively. **(C)** Alphafold model of the RING-like domain of Sne_A19470 shows a divergent RING fold (blue) where only the ligands of the Zn_2_ ion (magenta) are conserved. **(D)** Multiple alignment of the RING-like region of Sne_A19470 with the two best DALI hits (pdb:8A38 and pdb:6YXE). Coloring as in B.

**Supplementary Figure 2. Immunoblot detection of ubiquitin-linked chains generated by SneRING. (A-E)** *In vitro* autoubiquitination assay as in Fig. 2A was performed for 5 h at 37 °C. The reaction was analyzed by SDS-PAGE and western blot, using primary antibodies against **(A)** K11-, **(B)** K63-, **(C)** K48-, **(D)** K6- **(C)** and **(E)** M1-linked ubiquitin chains. Positive controls were either purchased (K6- and K48-linked chains in C and D, respectively) or synthesized *in vitro* (M1-linked ubiquitin chains, E).

**Supplementary Figure 3. SneRING is expressed during infection in different host cells.** **(A, B)** HeLa229 or U2OS cells were infected with Sne at an MOI 1 and mRNA was isolated on day two and four pi. **(C)** THP-1 cells were differentiated into macrophage-like cells using PMA, infected with Sne, and mRNA was isolated on day two and four pi. **(D)** Primary human “M2”-like macrophages were derived from peripheral blood monocytes (M-CSF/IL-4). After infection with Sne, mRNA was isolated on day two, four, and six pi. **(E)** *A. castellanii* was infected with Sne and four days pi, mRNA was isolated. A modified 2^-ΔΔCt^ method was used to quantify the expression of SneRING and SnGroEL for each time point. ΔCt values were calculated relative to the human reference gene YWHAZ (A, B, C, and D) or 5S RNA of *A. castellanii* (E). log_2_ fold change (log_2_FC/2^-ΔΔCt^) for each tested Sne gene vs the non-infected control samples was calculated. Expression levels relative to Sne 5S RNA are shown as mean ± SD from independent biological replicates.

**Supplementary Figure 4.** **SneRING distribution in cells after expression.** **(A)** A human codon-adjusted, FLAG-tagged version of the SneRING was transfected in HeLa229 cells expressing mitochondria-targeted GFP (Mito-GFP, green channel). 24 h after transfection, the cells were either left uninfected (-Sne) or were infected with Sne at an MOI 1 (+Sne). On day 3 pi, the cells were fixed and stained with DAPI (blue channel), and primary antibodies against the FLAG-tag (red channel) and SnGroEL (magenta channel), followed by a fluorophore-coupled secondary antibody. **(B)** HeLa229 cells stably expressing ER-targeted KDEL-dsRED (red channel) were transfected with a human codon-adjusted FLAG-tagged version of the SneRING. 24 h post-transfection, one set of samples was infected with Sne for 3 days (+Sne), when they were fixed together with control, non-infected cells (-Sne) and stained using DAPI (blue channel), and antibodies against the FLAG-tag (green channel) and SnGroEL (magenta channel), followed by staining with fluorophore-coupled secondary antibodies. All images were taken using laser confocal scanning microscopy. The scale bar is 10 µm.

**Supplementary Figure 5. Potential *S. negevensis* proteins that interact with SneRING.** The graph shows identified bacterial proteins from samples described in Fig 5A, with significance (-log_10_p-value) plotted against the log_2_ fold change (log_2_FC) of transfected/infected U2OS cells (U2OS+Sne+SneRING) relative to non-transfected/infected controls (U2OS+Sne) Enriched proteins are presented in blue, reduced proteins are labeled in red, and unchanged proteins are grey.

**Supplementary Table 1.** **List of significantly enriched host cell and bacterial potential interactors of SneRING.** Significantly enriched proteins in pull-down samples described in Fig 5 and Fig S5 with log_2_Fold Change > 1 are listed in descending order. Darker color shade represents higher values. SneRING (SNE_A12920) is highlighted in yellow.
