## Supplemental Figure 1 for "Identification and characterization of a ubiquitin E3 RING ligase of the *Chlamydia*-like bacterium *Simkania negevensis*"

Figure S1

**A**

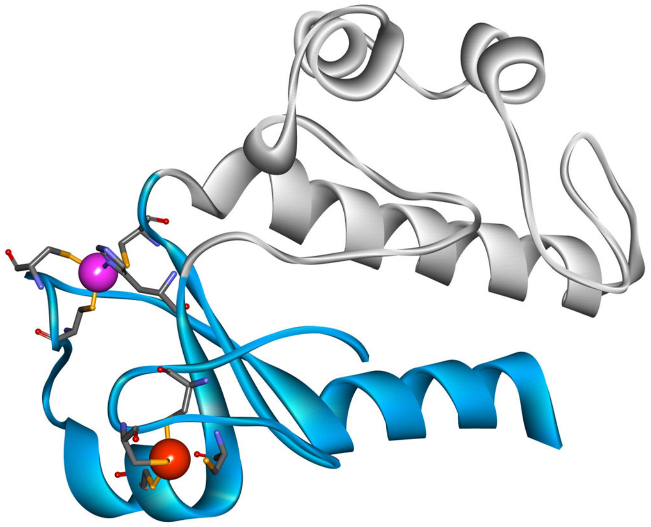


**B**

```
Sne A08700      VNFNCSICLD........SVSKKFAALDCNFGKERLALIDKILDEQGHAWKEGRQLP.W.IDDWMRID........PKKL
Sne A08690      DNLECAISLE........KPMDHL.VLGCNEGIERVRVIDEMA....KAIAESDIRPTW.LRHNGSVE........VSKL
Chl-KDK64       LPLFCHLTQESYCGEQPNPVIPAMVFFDCNFAKERLALMDQIVLSG............WYQEHLGRMT.......LETV
Ver-JSS10       GIPDCPIGQS........PIANPL.FADCNEASVRVDLMDKICQAIFEARKNTQSDPAWYVDNKIRFDEAMTALNIPPEI
Neptunochl.     TPLQCPFSLEKYTGGNPNPELTAMAFSDCNFGLERLELMDRILSS............EWYVFHQGRFK........MGEI
5DKA (RNF125)   TSFDCAVCLE.........VLHQPVRTRC...................................................

Sne A08700      EKAYFITAEYLPLLDPQQDESKKLLTNEGYFNLAGLHGGCKECYDSVITRK......CECPICRAQHVQVYTSKY
Sne A08690      RETITMSDMF..........AKKLLTEEGFVKLSDLHFFDRESVNGSTQRSGMERDYTKCDKCRAELEEVTTSKA
Chl-KDK64       RERFGMSDQLFRE....VQSSKCLLSSDGKLNIEGLHGGERRSMSMRRTYS......QDCPVCRGRG.QLVHSVA
Ver-JSS10       RNKIGPHAPL..........FKRLLTAEGVFNKDGLHIYERENLINWVYVN....LHGRCPECTQPS.RTSFSKA
Neptunochl.     KVEFGVTNALFNA....ALGSEKLLSSDGLFNLDGLHGGERRTIQKHVSYS......NNCPMCHNRVRGLYQSRY
5DKA (RNF125)   ...................................GHVFCRSCIATSLKNN.....KWTCPYCRAYLPSEGVPAT
```

**C**

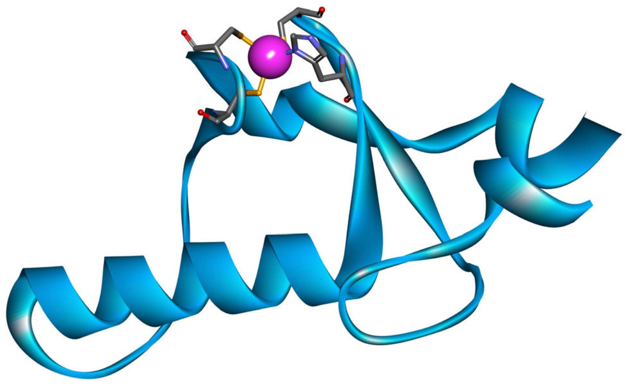


**D**

```
Sne A19470      IANLQDPISLERTRNPVISEKCFHTFQRSGISGTLYTDLESRLGNWDLRVPHEYECALCKEKTPIEQIVTN
8A38 (TRIM2)    KQFLICSICLERYKNPKV.LPCLHTFCERCLQNYIPAH............SLTLSCPVCRQTSILPEKVAA
6YXE (TRIM69)   TMELHCPLCNDWFRDPLM.LCSGHNFCEACIQDFWRLQ............AKETFCPECKMLCQYNNCTFN
```
