## Supplementary figures and images for "Identification and characterization of a ubiquitin E3 RING ligase of the *Chlamydia*-like bacterium *Simkania negevensis*"

### Supplemental Figure 2

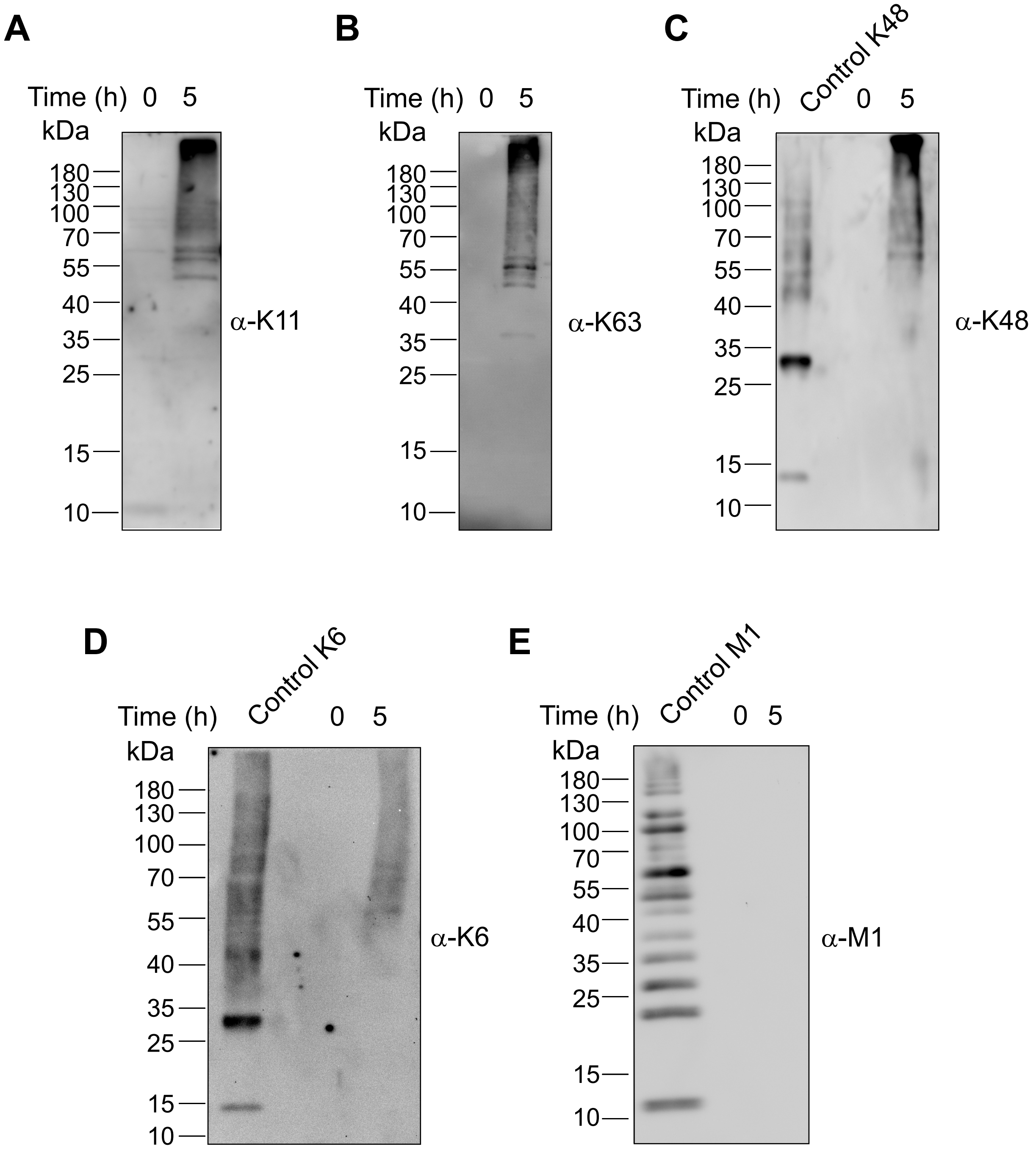

### Supplemental Figure 3

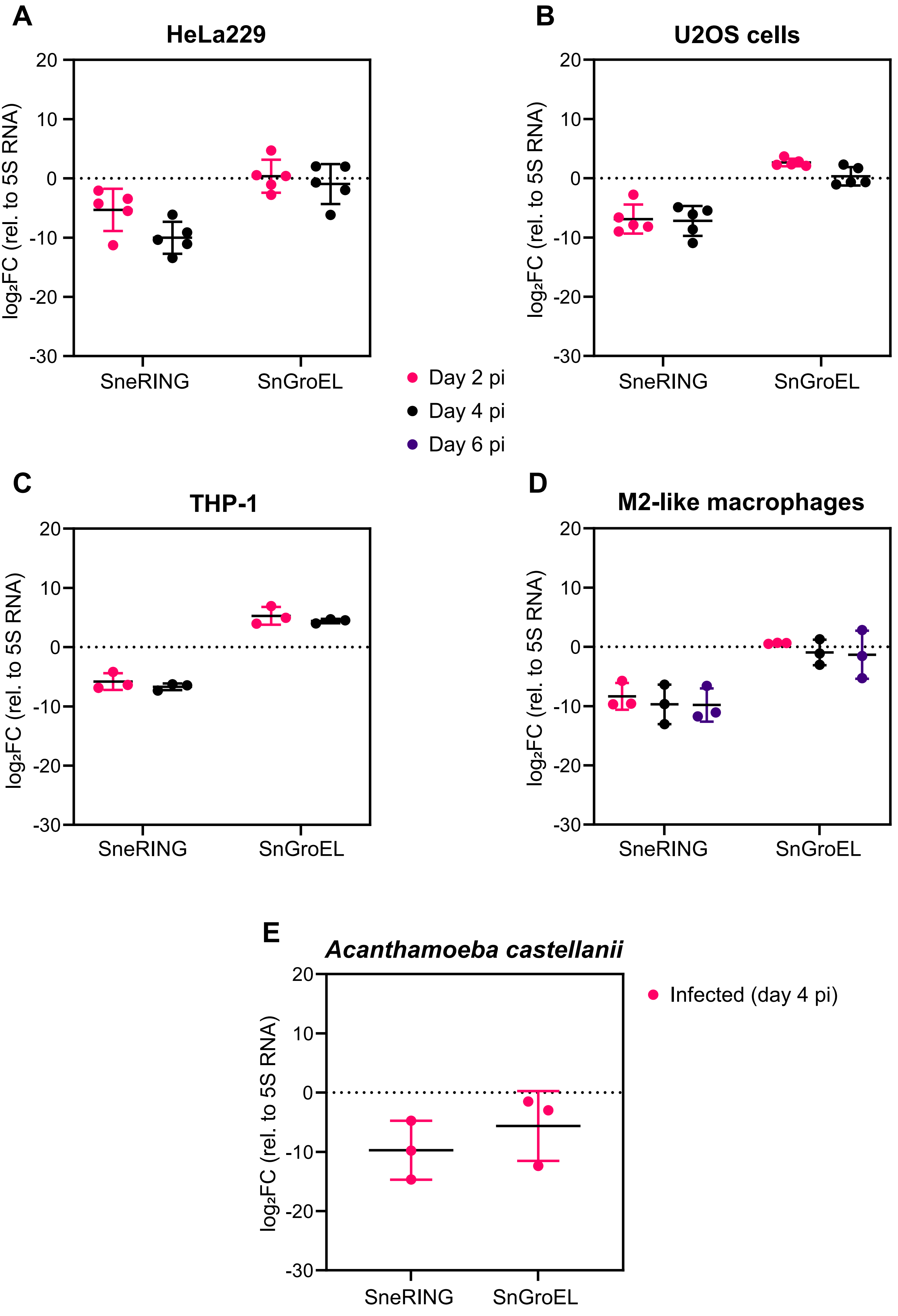

### Supplemental Figure 4

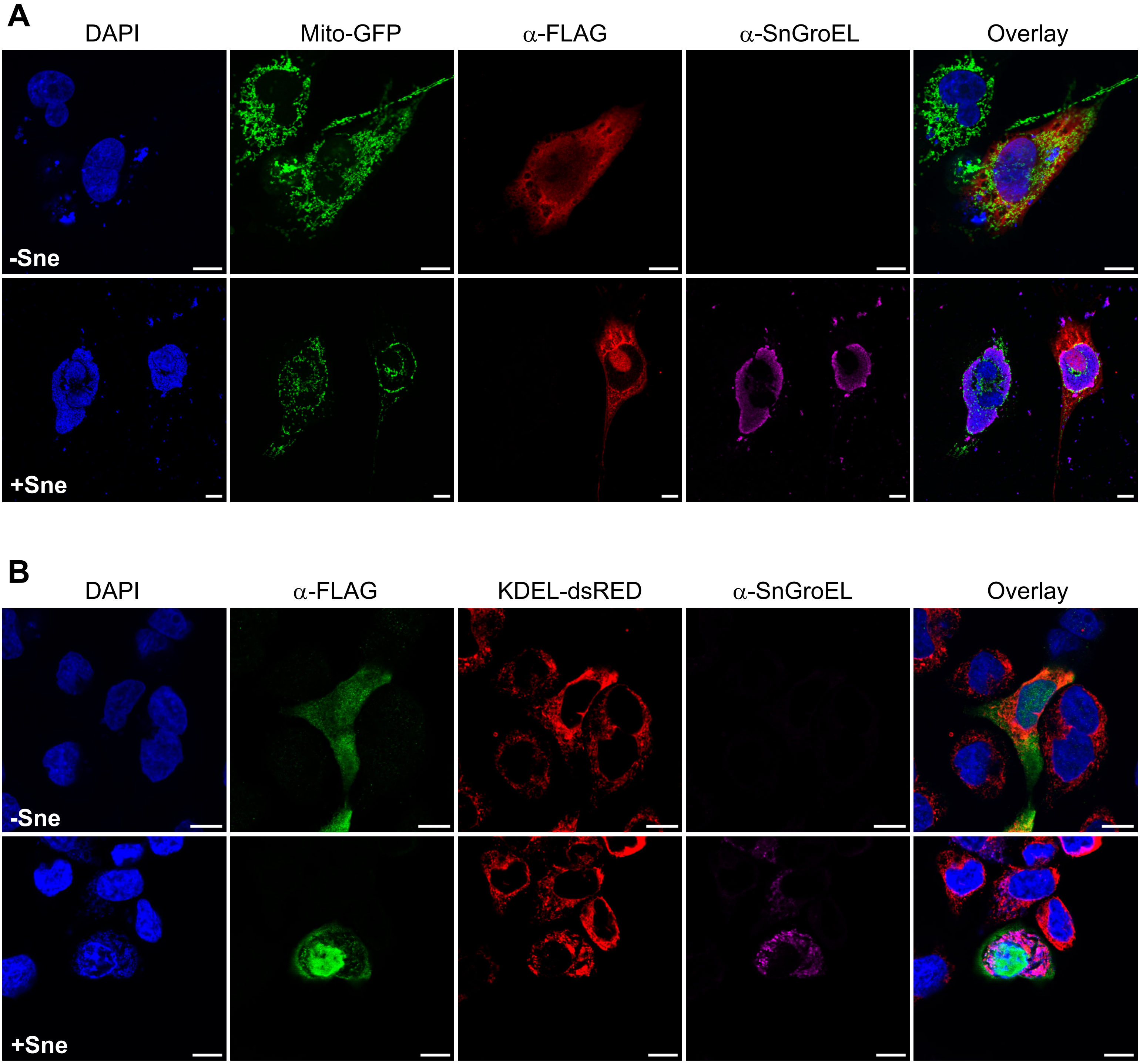

### Supplemental Figure 5

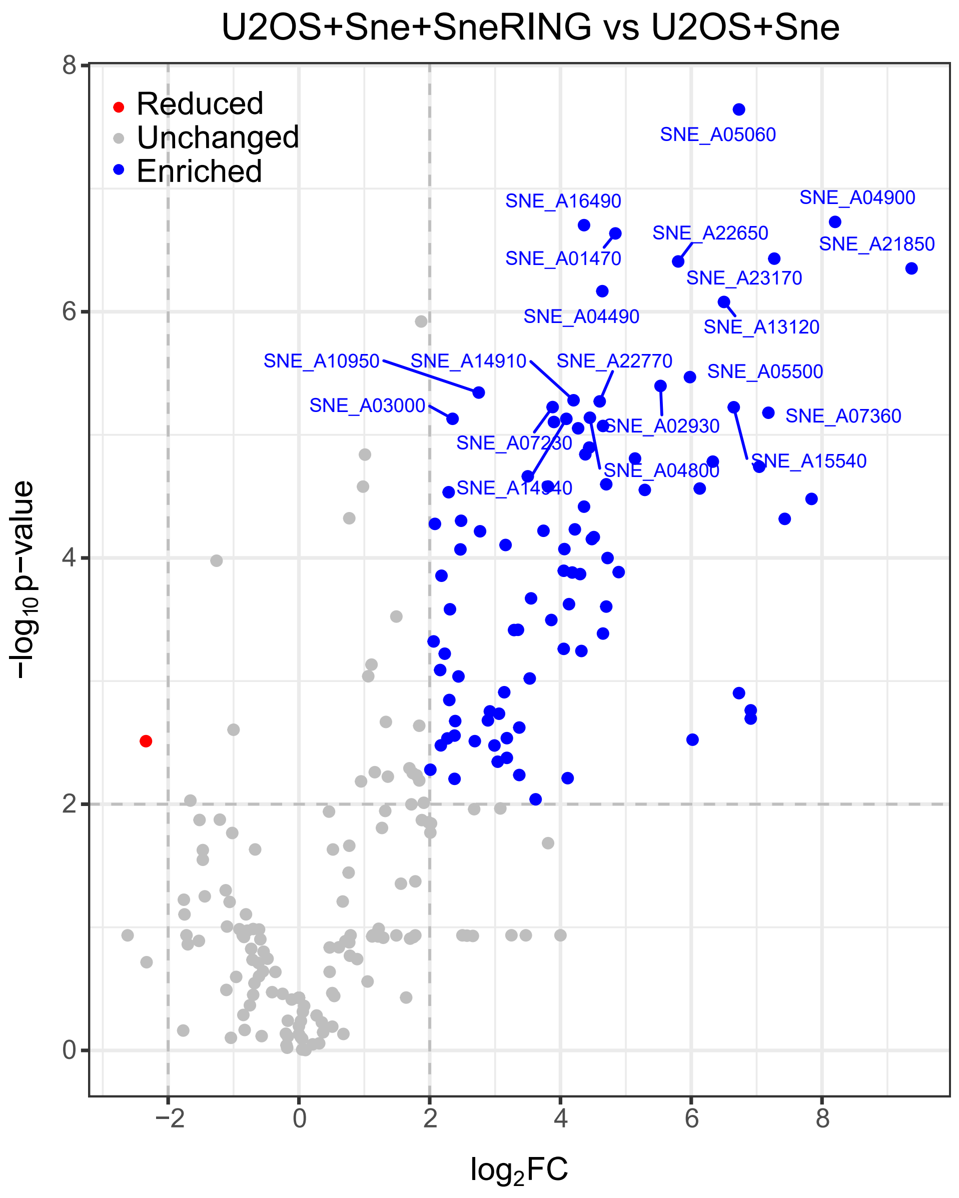
